# Structure and design of Langya virus glycoprotein antigens

**DOI:** 10.1101/2023.08.20.554025

**Authors:** Zhaoqian Wang, Matthew McCallum, Lianying Yan, William Sharkey, Young-Jun Park, Ha V. Dang, Moushimi Amaya, Ashley Person, Christopher C. Broder, David Veesler

**Author notes:** These authors contributed equally (Zhaoqian Wang, Matthew McCallum, Lianying Yan).

## Abstract

Langya virus (LayV) is a recently discovered henipavirus (HNV), isolated from febrile patients in China. HNV entry into host cells is mediated by the attachment (G) and fusion (F) glycoproteins which are the main targets of neutralizing antibodies. We show here that the LayV F and G glycoproteins promote membrane fusion with human, mouse and hamster target cells using a different, yet unknown, receptor than NiV and HeV and that NiV-and HeV-elicited monoclonal and polyclonal antibodies do not cross-react with LayV F and G. We determined cryo-electron microscopy structures of LayV F, in the prefusion and postfusion states, and of LayV G, revealing previously unknown conformational landscapes and their distinct antigenicity relative to NiV and HeV. We computationally designed stabilized LayV G constructs and demonstrate the generalizability of an HNV F prefusion-stabilization strategy. Our data will support the development of vaccines and therapeutics against LayV and closely related HNVs.

---

Nipah virus (NiV) and Hendra virus (HeV) are bat-borne zoonotic pathogens of the Henipavirus (HNV) genus causing severe encephalitis and respiratory symptoms in humans, with fatality rates between 50 and 100%^1^. NiV has spilled over to humans on a regular basis in Bangladesh as well as in India and the detection of HNVs in other continents underscore their global distribution^2–7^. Moreover, the recent discovery of a previously unknown HeV genotype in Australia is an urgent reminder of the pandemic threat HNVs represent^8,9^. To date, no approved vaccines or therapeutics exist for use in humans against any HNVs^10^.

HNV infections are facilitated by two viral surface glycoproteins, known as attachment glycoprotein (G) and fusion glycoprotein (F), coupling receptor engagement to fusion of the viral and host membranes. The homotetrameric type II transmembrane protein G is the receptor-binding protein and comprises an N-terminal cytoplasmic domain, a transmembrane domain, a stalk domain, a neck domain and a C-terminal receptor binding head domain^11^. The homotrimeric type I transmembrane protein F is the viral fusion protein which is cleaved into disulfide-linked F_1_ and F_2_ subunits by cathepsin L through an endosomal recycling process^12–14^. Upon G-mediated receptor engagement, F is triggered and undergoes large-scale conformational changes promoting membrane fusion and viral entry^15^. F and G are the targets of neutralizing antibodies which are correlates of protection against NiV and HeV^16–23^.

NiV, HeV, Ghanaian virus (GhV) and Cedar virus (CedV) utilize a subset of ephrin receptors to gain entry into host cells^24–28^. Furthermore, although most HNVs are bat-borne viruses, including the recently discovered Angavokely virus (AngV)^7^, Mojiang virus (MojV) was discovered in rats^29^ whereas Gamak (GAKV) and Daeryong (DARV) HNVs were identified in shrews^30^. Most recently, Langya virus (LayV) was found in shrews and also isolated from febrile patients in China^31^, thereby representing the first confirmed human-infecting HNV from shrew origin. Phylogenetic analyzes indicate that LayV, MojV, GAKV and DARV cluster together and are distantly related to bat-borne HNVs^30–32^. Although receptor usage for MojV is known to differ from that of NiV, HeV, GhV and CedV, the specific host receptor(s) used by MojV, LayV, GAKV and DARV remain elusive^32^.

The LayV F and G glycoproteins are most closely related to MojV F and G, sharing 90% and 86% respective amino acid sequence identity, but distantly related to the better studied NiV and HeV counterparts. As a result, it is expected that countermeasures in development for NiV and HeV will not be effective against these divergent viruses^32^, including the newly emerged human-infecting LayV. The scarcity of structural and functional information for the LayV and MojV F and G glycoproteins combined with their weak sequence relatedness to NiV/HeV F and G limit our ability to assess conserved and divergent architectural traits for the development of countermeasures and antigen design strategy broadly applicable to the HNV genus^32–34^. To address this knowledge gap, we determined cryo-electron microscopy structures of LayV F in the prefusion and postfusion conformations, revealing the previously uncharacterized refolding events underlying HNV F-mediated membrane fusion. Although NiV/HeV-elicited polyclonal and monoclonal antibodies (mAbs) did not cross-react with LayV F and G, we identified a MojV F-directed monoclonal antibody (mAb) (4G5) binding to LayV F, and unveiled the molecular basis of recognition, as well as a MojV G-directed mAb (2B2) cross-reacting with LayV G. We determined a cryoEM structure of LayV G, which adopts a distinct conformation than that previously described for NiV G or other paramyxovirus attachment proteins, defining another snapshot of the host invasion process. We computationally designed stabilized LayV G constructs and demonstrate the broad applicability of an HNV F prefusion-stabilization strategy which will pave the way for vaccine development efforts. Collectively, our data provide molecular blueprints of the two key targets of LayV vaccine design and pave the way for designing countermeasures against this emerging pathogen.

## Functional assessment of the LayV F and G glycoproteins

LayV F possesses a putative cleavage site at residue 104 (R104), as is the case for NiV (R109) and HeV (K109), but lacks the YSRL and YY motifs present in the NiV/HeV F C-terminal cytoplasmic domains to promote endosomal recycling and subsequent cathepsin L-mediated F cleavage^12,35^ (Fig. **1a**). Nevertheless, we observed that transient transfection of LayV F or LayV F and G in CHO-K1, HEK293T or mouse Neuro-2a cells resulted in production of the F_0_ precursor and proteolytically cleaved F_1_ (and F_2_) with approximately similar ratio to that observed for other HNVs (Fig. **1b-d**). These data are reminiscent of findings made with MojV F^36^ and suggest that the LayV/MojV F proteolytic cleavage pathway is likely distinct from that described for NiV and HeV F.

**Figure 1:**
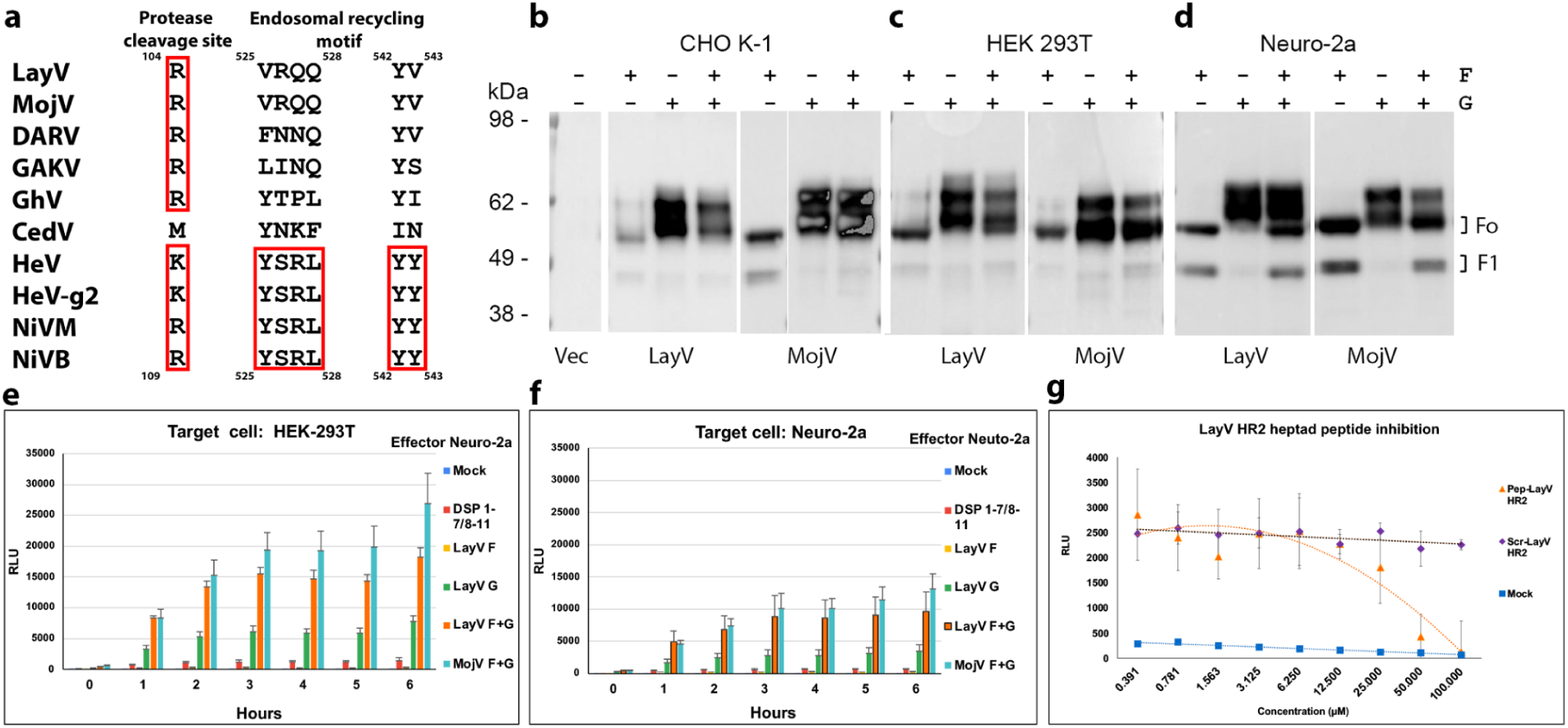
Functional assessment of the LayV F and G glycoproteins. a, HNV F sequence alignment focused on the activating protease cleavage site and the endosomal recycling motif. Red rectangles highlight conserved/conservatively substituted residues/motifs. LayV F sequence numbering is shown on top whereas NiV F sequence numbering is shown at the bottom. HeV-g2 refers to a recently discovered HeV variant; NiVM refers to NiV-Malaysia; NiVB refers to NiV-Bangladesh. **b-d,** Assessment of HNV F cleavage in CHO-K1 (b), HEK293T (c) or mouse Neuro-2a cells (d) using Western blot with rabbit anti-S peptide:HRP antibodies. Transfected plasmids are indicated above each lane with “F” and “G” labels. **e-f,** Effector Neuro-2a cells were transfected with LayV F, LayV G, LayV F and LayV G, MojV F, MojV G or pcDNA3.1-MojV F and MojV G along with the DSP 1-7 plasmid. Target (HEK293T or Neuro-2a) cells were transfected with DSP 8-11 to enable luciferase detection upon cell/cell fusion. Mock: cells transfected with an empty pcDNA3.1 plasmid **g,** Concentration-dependent inhibition of cell-cell fusion in presence of either a LayV F HR2 peptide or a scrambled peptide of identical composition. Effector Neuro-2a cells were co-transfected with LayV F, LayV G and a DSP 1-7 plasmid. Target HEK293T cells were transfected with DSP 8-11.

We subsequently assayed the ability of LayV F and G to promote cell-cell fusion using a split luciferase system. Transient transfection of Neuro-2a effector cells with LayV F and G resulted in detectable fusion with target HEK293T or Neuro-2a cells (Fig. **1e-f**). Furthermore, a LayV F HR2 peptide, but not a scrambled peptide with identical composition, inhibited LayV F/G-mediated fusion in a concentration-dependent manner, demonstrating that the observed membrane fusion was promoted by F/G (Fig. **1g**). These data show that the LayV F and G glycoproteins are fusion-competent although they promote cell-cell membrane fusion much more weakly than their NiV/HeV counterparts, as was the case for MojV F and G^32,36^.

### Structures of LayV F in the prefusion and postfusion conformations

To understand the architecture of LayV F, we recombinantly produced a LayV F glycoprotein ectodomain construct C-terminally-fused to a GCN4 trimerization motif^14,16,17^. Using electron microscopy (EM) imaging of negatively stained samples, we observed that LayV F formed well-folded compact homotrimers, which are typical of the prefusion state, but that the protein spontaneously refolded to the postfusion state over time (Extended Data Fig. 1). As a result, we collected two cryoEM datasets 4 months apart post purification and determined structures of prefusion F and postfusion F at 2.7 Å and 3.5 Å resolution, respectively (Extended Data Fig. 2 and Table **1**).

**Table 1.** CryoEM data collection and refinement statistics.

|  | LayV G<br>(EMDB<br>41643)<br>(PDB<br>8TVI) | LayV F<br>prefusion<br>(EMDB<br>41640)<br>(PDB<br>8TVF) | LayV F<br>postfusio<br>n<br>(EMDB<br>41639)<br>(PDB<br>8TVE) | 4G5-bou<br>nd LayV<br>F (global<br>refineme<br>nt)<br>(EMDB<br>41644) | 4G5-bo<br>und LayV F<br>(local<br>refineme<br>nt<br>proxim<br>al)<br>(EMDB<br>41642)<br>(PDB<br>8TVH) | LayV F<br>4G5<br>(local<br>refineme<br>nt<br>distal)<br>(EMDB<br>41641)<br>(PDB<br>8TVG) | GhV F<br>(EMDB<br>41636)<br>(PDB<br>8TVB) |
| --- | --- | --- | --- | --- | --- | --- | --- |
| **Data collection and processing** |  |  |  |  |  |  |  |
| Magnification | 45,000 | 105,000 | 45,000 | 105,000 | 105,000 | 105,000 | 105,000 |
| Voltage (kV) | 200 | 300 | 200 | 300 | 300 | 300 | 300 |
| Electron exposure<br>(e-/Å<sup>2</sup>) | 47 | 63 | 47 | 63 | 63 | 63 | 63 |
| Defocus range (µm) | 0.5 - 2.4 | 0.7 -1.7 | 0.7 -1.7 | 1.3 -1.7 | 1.3 -1.7 | 1.3 -1.7 | 0.5 -2.5 |
| Pixel size (Å) | 0.89 | 0.843 | 0.89 | 0.843 | 0.843 | 0.843 | 0.843 |
| Symmetry imposed | C4 | C3 | C3 | C3 | C3 | C3 | C3 |
| Initial particle<br>images (no.) | 1,374,036 | 295,258 | 681,389 | 283,620 | 283,620 | 283,620 | 349,279 |
| Final particle<br>images (no.) | 145,167 | 53,917 | 86,749 | 60,440 | 60,440 | 60,440 | 60,207 |
| Map resolution (Å)<br>FSC threshold | 3.2<br>0.143 | 2.8<br>0.143 | 3.9<br>0.143 | 3.5<br>0.143 | 3.6<br>0.143 | 3.3<br>0.143 | 2.9<br>0.143 |
| **Refinement** |  |  |  |  |  |  |  |
| Model resolution (Å)<br>FSC threshold | 3.4<br>0.5 | 3.0<br>0.5 | 4.0<br>0.5 |  | 3.7<br>0.5 | 3.6<br>0.5 | 3.1<br>0.5 |
| Model resolution<br>range (Å) |  |  |  |  |  |  |  |
| Map sharpening *B*<br>factor (Å<sup>2</sup>) | -297 | -97 | -137 | -118 | -96 | -112 | -108 |
| Model composition<br>Non-hydrogen<br>atoms<br>Protein residues<br>Ligands | 16,120<br>2,056<br>8 | 11,376<br>1,296<br>12 | 9,204<br>1,233<br>6 |  | 6,915<br>948<br>3 | 8,583<br>1,131<br>6 | 9,894<br>1,266<br>12 |
| *B* factors (Å<sup>2</sup>)<br>Protein<br>Ligand | 23.19<br>41.31 | 14.17<br>26.09 | 29.12<br>45.84 |  | 29.44<br>32.00 | 23.17<br>33.09 | 14.69<br>19.75 |
| R.m.s. deviations |  |  |  |  |  |  |  |
| Bond lengths (Å) | 0.01 | 0.01 | 0.01 |  | 0.01 | 0.01 | 0.01 |
| Bond angles (°) | 0.94 | 1.29 | 1.43 |  | 1.22 | 1.23 | 0.91 |
| Validation |  |  |  |  |  |  |  |
| MolProbity score | 1.01 | 0.56 | 1.20 |  | 0.91 | 0.89 | 0.73 |
| Clashscore | 1.63 | 0.15 | 3.58 |  | 0.98 | 0.70 | 0.70 |
| Poor rotamers (%) | 0.00 | 0.00 | 0.32 |  | 0.48 | 0.65 | 0.00 |
| Ramachandran plot |  |  |  |  |  |  |  |
| Favored (%) | 97.54 | 98.83 | 97.79 |  | 97.40 | 97.05 | 98.56 |
| Allowed (%) | 2.26 | 0.93 | 2.21 |  | 1.95 | 2.68 | 1.20 |
| Disallowed (%) | 0.20 | 0.23 | 0.00 |  | 0.65 | 0.27 | 0.24 |

Prefusion LayV F folds as a ∼90Å-high and ∼90Å-wide pyramidal-shaped trimer, typical of HNV F glycoproteins in the prefusion conformation (Fig. 2a**-b**). Although LayV F shares only 44% amino acid sequence identity with NiV F, these two glycoproteins adopt a strikingly similar tertiary and quaternary structures^13,14,16,17^ (Fig. 2a**-d**). Indeed, a LayV F protomer can be superimposed to NiV F with root mean square deviation of 2.3 Å over 432 aligned Cα pairs as compared to a root mean square deviation of 1.1 Å over 436 aligned Cα pairs for NiV F and HeV F (Fig. 2c). All 5 disulfide bonds within a single LayV F protomer are fully conserved compared with NiV/HeV F, suggesting they participate in proper HNV F folding. The LayV F fusion peptide sequence (residues 110-122) is identical to the MojV F fusion peptide and highly conserved with NiV/HeV F and they all exhibit an identical structural organization in the prefusion trimers (Fig. 2e**-h**). Both N-linked glycans present in LayV F are resolved in the cryoEM map (at position N65 and N459), as compared to NiV and HeV F which harbor at least four N-linked oligosaccharides^37,38^. The LayV F N65 glycan protrudes from the trimer apex from a roughly similar location to the NiV/HeV N67-linked glycan (Fig. 2a**-c**) which is part of an antigenic site targeted by several NiV F-neutralizing antibodies^16,39^. Based on the marked sequence divergence and distinct glycosylation profiles of LayV F relative to NiV and HeV F, it is expected that LayV F will have a markedly distinct antigenicity.

**Figure 2:**
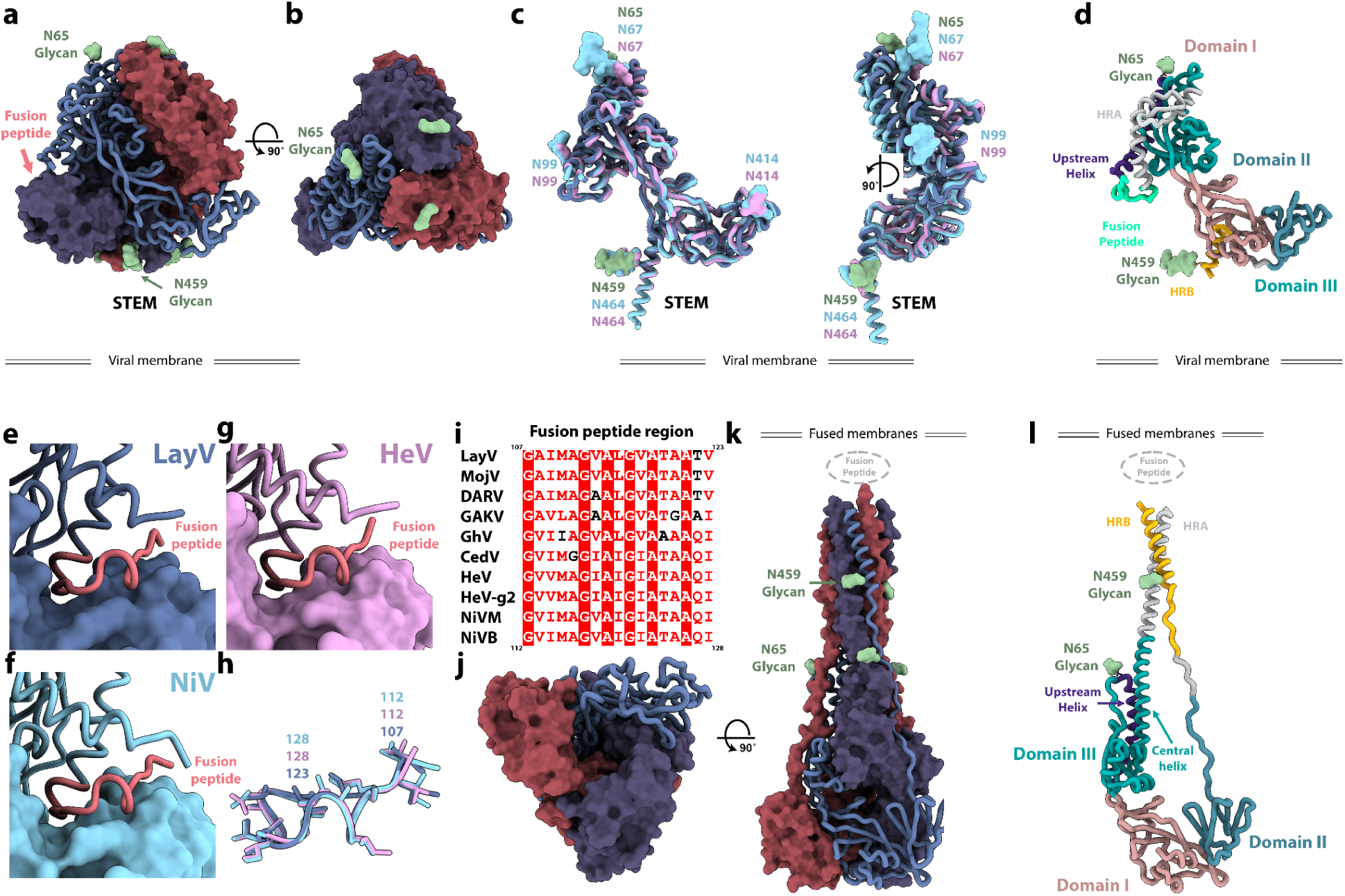
Structures of LayV F in the prefusion and postfusion conformations. a-b, Prefusion LayV F trimer structure in two orthogonal orientations. Each protomer is colored distinctly and one of them is shown as ribbons whereas the other two protomers are rendered as surfaces. N-linked glycans are shown as green surfaces and labeled. **c,** Superimposition of LayV F (dark blue) with NiV F (cyan, PDB 7KI4) and HeV F (pink, PDB 7KI6). N-linked glycans are rendered with the same color scheme and labeled. **d,** Ribbon diagram of a LayV F protomer in the prefusion state colored and annotated by functional domains^17^. **e-g** Zoomed-in views of LayV F (e, blue), NiV F (f, cyan) and HeV F (g, pink) trimers focused on the fusion peptides (salmon, ribbon view) interacting with the neighboring protomer (rendered as a surface). **h,** Superimposition of LayV F (blue), NiV F (cyan) and HeV F (pink) fusion peptides with side chains shown as sticks. **i,** Sequence alignment focused on the HNV F fusion peptide region. Color scheme matches ESPript: red background with white text indicates identical region; white background with red text indicates conserved region; white background with black text indicates outliers. LayV F sequence numbering is shown on top, while NiV F sequence number is shown at the bottom. HeV-g2 refers to a recently discovered HeV variant; NiVM refers to NiV-Malaysia; NiVB refers to NiV-Bangladesh. **j-k,** Postfusion LayV F structure in two orthogonal orientations with the same color scheme as panels a-b. N-linked glycans are rendered as green surfaces. **l,** Ribbon diagram of a LayV F protomer in the postfusion state colored and annotated by functional domains.

Postfusion LayV F forms a ∼150 Å-high and ∼70 Å-wide conical trimeric structure with a prominent triple helical bundle assembled from the central helices and HR1, surrounded by three antiparallel HR2 helices yielding a six-helix bundle at one end of the trimer. The opposite end of the molecule forms a triangular base (Fig. 2j**-l**). The HR1 and the HR2 motif of each F protomer interacts exclusively with the other two protomers within a trimer forming an intertwined network of contacts (Fig. 2j**-l**). The cryoEM map resolves glycans at position N65 and N459 decorating the periphery of the elongated trimer (Fig. 2). In contrast to prefusion F, the fusion peptide and transmembrane regions would be located at the same end of the cone-like structure to promote fusion of the viral and host membranes. Postfusion LayV F shares its overall topology with other paramyxovirus postfusion F trimers as well as with postfusion coronavirus S trimers (Extended Data Fig. 3), highlighting the evolutionary relationships among the fusion machinerysu of these distinct viruses^40–45^.

Although F refolding affects a large fraction of the trimer, the N-terminus, the β-rich domains comprising residues 281-420, and the upstream helix remain mostly unchanged, besides modification of their relative orientations, as is the case for the disulfide bonds (Fig. 2d**,l**). This conformational transition leads to an approximately four-fold enhancement of the surface area buried at the interface between each pair of protomers within the prefusion F state (∼2,250 Å^2^) and the postfusion F state (∼9,000 Å^2^), underscoring the irreversible nature of these structural rearrangements^46^.

### A generalizable prefusion stabilization strategy for HNV F glycoproteins

Spontaneous refolding of the recombinant LayV F ectodomain trimer from the prefusion state to the postfusion conformation underscores the metastability of this glycoprotein, as is the case for numerous other viral fusion proteins^40–42,45–50^. Previous studies showed that immunization with prefusion NiV F or HeV F, but not postfusion F, allowed elicitation of neutralizing antibodies against these vaccine targets^46,51^, thereby motivating the identification of suitable strategies to stabilize F in the prefusion state. Hence, we evaluated the portability of several previously described NiV/HeV prefusion-stabilizing mutations to the LayV F trimer. We evaluated (i) the NiV L172F (cavity-filling, LayV F I167F) and S191P (postfusion central helix breaker, LayV F S186P) substitutions previously identified for NiV F^51^ and (ii) the engineered disulfide bond spanning the F_2_ and F_1_ subunits (NiV/HeV N100C/A119C corresponding to LayV F N95C/A114C), near the proteolytic F cleavage site, originally described for HeV F^13^ and subsequently ported to NiV F^17^, which appears to be compatible with our structure (Fig. 3a**-c)**. We observed that the I167F/S186P LayV F mutant mostly yielded postfusion trimers (Fig. 3e**-f**) whereas the LayV F N95C/A114C mutant enabled production of well-folded prefusion F trimers (Fig. 3g **and S1**). Combining all four mutations, however, produced prefusion F trimers along with some aggregates (Fig. 3h). These results indicate that the disulfide bond stapling the F_2_ and F_1_ subunits successfully reduced the LayV F metastability, stabilized the prefusion trimer and even rescued constructs yielding postfusion F.

**Figure 3:**
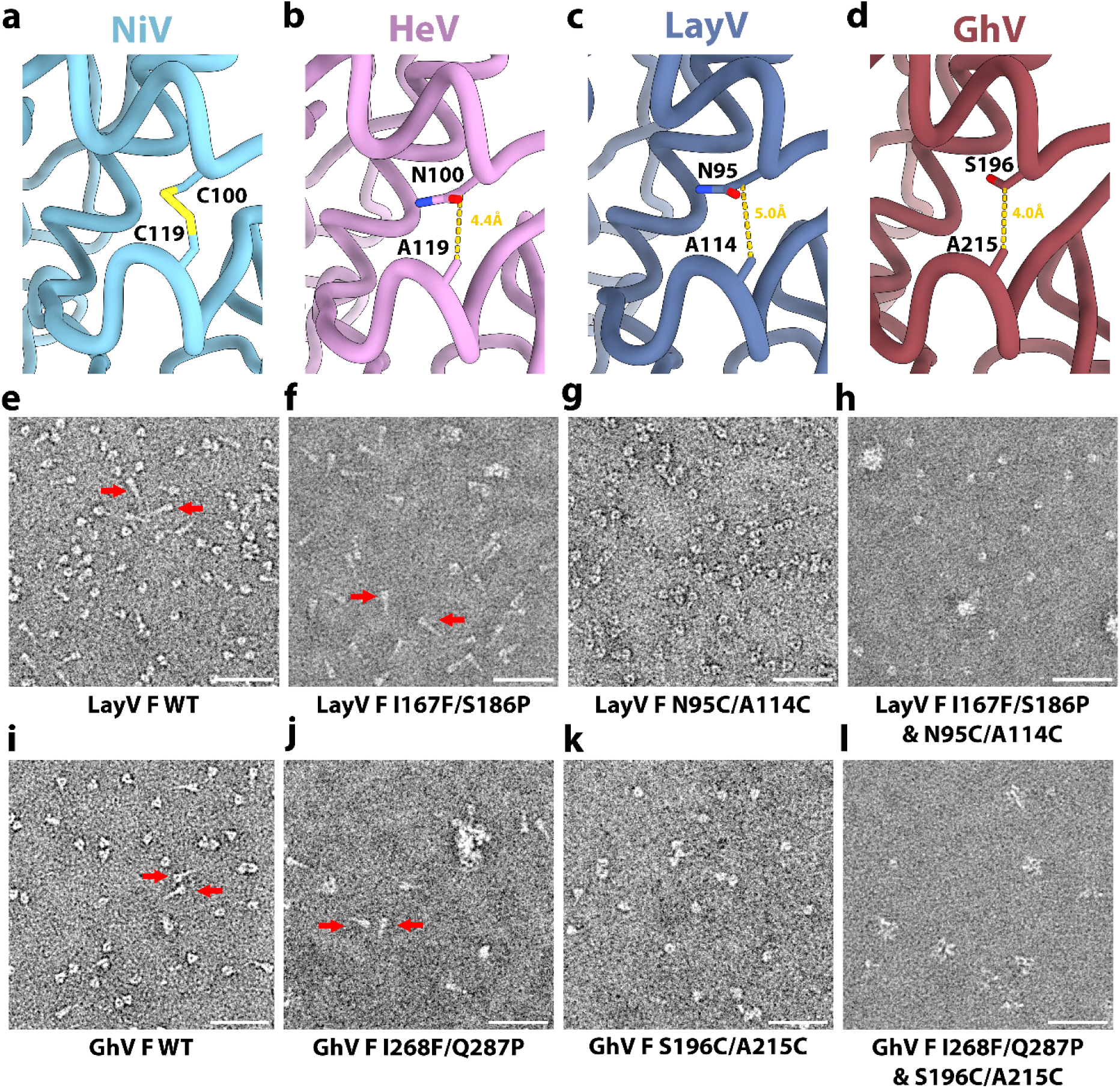
A generalizable prefusion stabilization strategy for HNV F glycoproteins. a-d, Zoomed-in views of NiV (PDB 6TYS) (a), HeV (PDB 7KI6) (b), LayV (c) and GhV (d) F structures highlighting the selected disulfide bond prefusion-stabilizing mutations. Yellow dashed lines show the distance between Cβ atoms of the labeled residues (corresponding to the mutated positions) suggesting possible compatibility with introduction of a disulfide. **e-h,** EM images of negatively stained, purified wildtype (e), I167F/S186P (f), N95C/A114C (g) and I167F/S186P/N95C/A114C (h) LayV F two days after purification. **i-l,** EM images of negatively stained, purified wildtype (i), I268F/Q287P (j), S196C/A215C (k) and I268F/Q287P/S196C/A215C (l) GhV F two days after purification. Scale bars: 50 nm. Red arrows indicate postfusion F trimers.

Based on these results, we decided to investigate if the same disulfide bond engineering strategy could be ported to GhV F, which shares 55% and 47% amino acid sequence identity with NiV F and with LayV F, respectively. To assist the design, we determined a cryoEM structure of prefusion GhV F at 2.9 Å resolution revealing a conserved overall HNV F architecture and suggesting that the S196C/A215C engineered disulfide bond would be compatible with our structure (Fig. 3a**-d** and Extended Data Fig. 4 and Table **1**). The I268F/Q287P and I268F/Q287P/S196C/A215C GhV F mutants mostly yielded aggregated and misfolded protein with a small fraction of postfusion trimers for I268F/Q287P and a small fraction of prefusion trimers for I268F/Q287P/S196C/A215C (Fig. 3j**-l**). Similar to LayV F, the GhV S196C/A215C F disulfide mutant led to enhanced expression of well-folded prefusion F trimer, relative to wildtype GhV F, underscoring that this disulfide stapling strategy is broadly generalizable to HNV F glycoproteins with marked sequence divergence (Extended Data Fig. 1).

### Identification of a LayV F cross-reactive monoclonal antibody

We previously showed that F-directed mAbs potently neutralize NiV and HeV, including (HeV-g2), and provide post-exposure protection against lethal challenge in ferrets^16–18,52^. We therefore evaluated the cross-reactivity of F-directed antibodies neutralizing NiV and HeV to the LayV F trimer immobilized at the surface of biolayer interferometry biosensors. Neither 5B3^17^ nor 12B2^16^ IgGs bound to LayV F (Fig. 4a), likely due to local structural differences (Extended Data Fig. 5). However, we observed robust binding of the 4G5 IgG, but not of the 3C4 IgG, to LayV F (Fig. 4a), two antibodies elicited through immunizations of mice with MojV F^36^. These results emphasize the close evolutionary and antigenic relationships between LayV F and MojV F, which share 90% amino acid sequence identity, along with their distant relatedness with the rest of the HNV genus.

**Figure 4:**
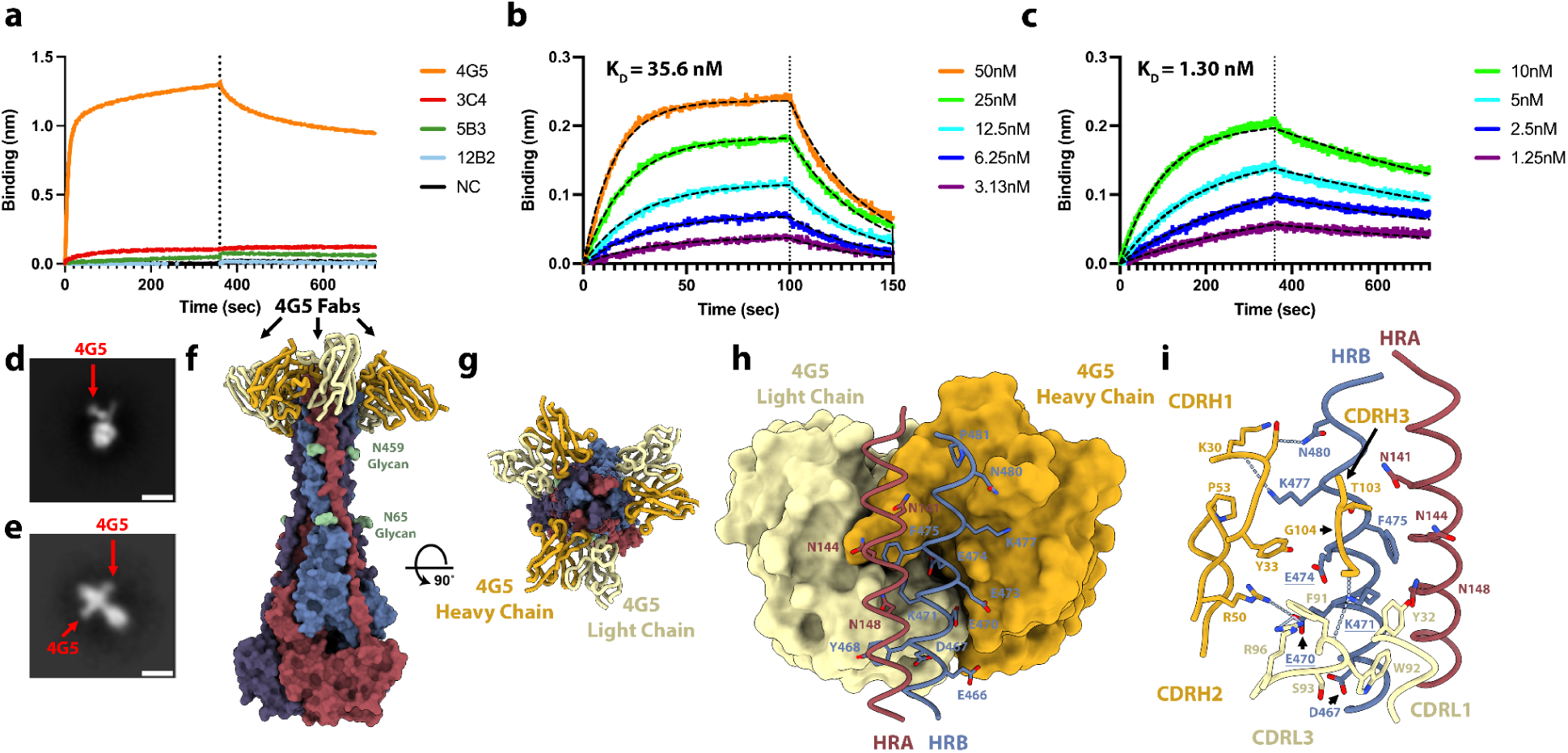
Identification of a LayV F cross-reactive monoclonal antibody. a, Evaluation of binding of F-directed mAbs at a concentration of 200nM to prefusion LayV N95C/A114C F immobilized at the surface of biolayer interferometry HIK1K biosensors. Negative control (black) is buffer only. **b-c,** Binding kinetics and affinity evaluation of 4G5 Fab binding to prefusion LayV N95C/A114C F (b) and postfusion (spontaneous refolded) wildtype LayV F (c) immobilized at the surface of biolayer interferometry HIS1K biosensors. The color key indicates the 4G5 Fab concentrations used and the black dashed lines show the fit to the data. **d-e,** Two dimensional EM class averages of negatively stained 4G5 bound to wildtype prefusion (d) and postfusion LayV F (e). Scale bar: 10 nm. **f-g,** 4G5-bound postfusion wildtype LayV F structure shown in two orthogonal orientations. Each protomer is colored distinctly and N-linked glycans are shown as green surfaces and labeled. 4G5 Fab heavy and light chains are rendered as gold and yellow ribbons, respectively. **h,** Zoomed-in view of the interactions between the 4G5 Fab (rendered as a surface colored gold and yellow for heavy and light chains, respectively) and LayV F HRA of one protomer (red) and HRB of the neighboring protomer (blue) with select epitope residues shown as sticks. **i,** Zoomed-in view of the interactions between the 4G5 Fab rendered as a surface (h) or ribbons (i) colored gold and yellow for heavy and light chains, respectively, and LayV F HRA of one protomer (red) and HRB of the neighboring protomer (blue). Selected epitope and paratope residues shown as sticks. Dashed lines indicate selected electrostatic interactions. Key epitope residue mutations between LayV F and NiV/HeV F are underlined.

To determine if 4G5 binding was specific for a given F conformation, we separately assessed binding to the prefusion-stabilized LayV F trimer (N95C/A114C) and to the postfusion F trimer (obtained through spontaneous refolding) using biolayer interferometry (Fig. 3h and Extended Data Fig.1). We found that the 4G5 Fab fragment recognizes prefusion and postfusion F with approximate affinities of 35.6 and 1.30 nM, respectively, with much slower off-rate for postfusion F than for prefusion F (Fig. 4b**-c**). We further validated these results using EM analysis of negatively stained samples revealing binding of one 4G5 Fab to each prefusion F trimer and of three Fabs to each postfusion F trimer (Fig. 4d**-e**).

To understand the molecular basis of 4G5 binding to LayV F, we determined a cryoEM structure of postfusion LayV F bound to the 4G5 Fab at 3.5 Å resolution (Fig. 4f**-g and S6**). We improved local map resolvability by carrying out local refinements of the 4G5-bound membrane-proximal region and of the membrane-distal conical base of the trimer yielding reconstructions at 3.6 and 3.3 Å, respectively (Fig. 4f**-g** and Extended Data Fig. 6 and Table **1**). 4G5 binds to an epitope located in the membrane proximal region of the postfusion F trimer forming an HRA/HRB 6-helix bundle. Both heavy and light chains contact HRA and HRB burying a surface of ∼700 Å^2^ at the interface with F using electrostatic interactions and shape complementarity (Fig. 4h**-i**). Specifically, the 4G5 epitope comprises HRA N141, N144 and N148 from one protomer and HRB E466, D467, Y468, E470, K471, E473, E474, F475, K477, G478 and N480 from a neighboring protomer. All these residues are strictly conserved between LayV and MojV F except for the N144S substitution, rationalizing the observed 4G5 cross-reactivity, whereas the numerous epitope substitutions (including the E470_LayV_/K475_NiV_, K471_LayV_/E476_NiV_ and E474_LayV_/R479_NiV_ charge reversals) likely account for the lack of 4G5 binding to NiV and HeV F (Fig. 4i and Extended Data Fig. 7). The tertiary structure of the HRA epitope moiety is partially rearranged whereas that of the HRB epitope moiety remains identical during the prefusion to postfusion F transition. However, these two regions are adjacent to each other in postfusion F but are spatially distant in prefusion F (Fig. 2). These observations explain the fact that we observed 4G5 bound to the prefusion F (HRB) stem, which accounts for a much larger fraction of contacts formed with the Fab than HRA, and that the affinity of 4G5 is greater for postfusion relative to prefusion F, as a result of more extensive interactions with postfusion F. Although 4G5 binds to both prefusion and postfusion F conformations, antibody binding might interfere sterically with the fusogenic conformational transition, as previously observed for the respiratory syncytial virus antibodies palivizumab and motavuzimab^42,53^.

### Computational design of a stabilized LayV G tetramer

NiV and HeV G are the targets of several potent neutralizing antibodies and the main focus of vaccine design against these pathogens. A HeV G vaccine is commercially available for use in horses in Australia (Equivac HeV, Zoetis Inc.), and a formulation suitable for use in humans was evaluated in phase 1 clinical trials^19–23,52,54,55^. To study this main target of the immune system in the newly emerged LayV, we recombinantly produced the LayV G ectodomain using transient transfection of Expi293 human cells. However, we could not detect expression of the wildtype LayV G ectodomain lacking the transmembrane and cytoplasmic domains (Fig. 5a).

**Figure 5.**
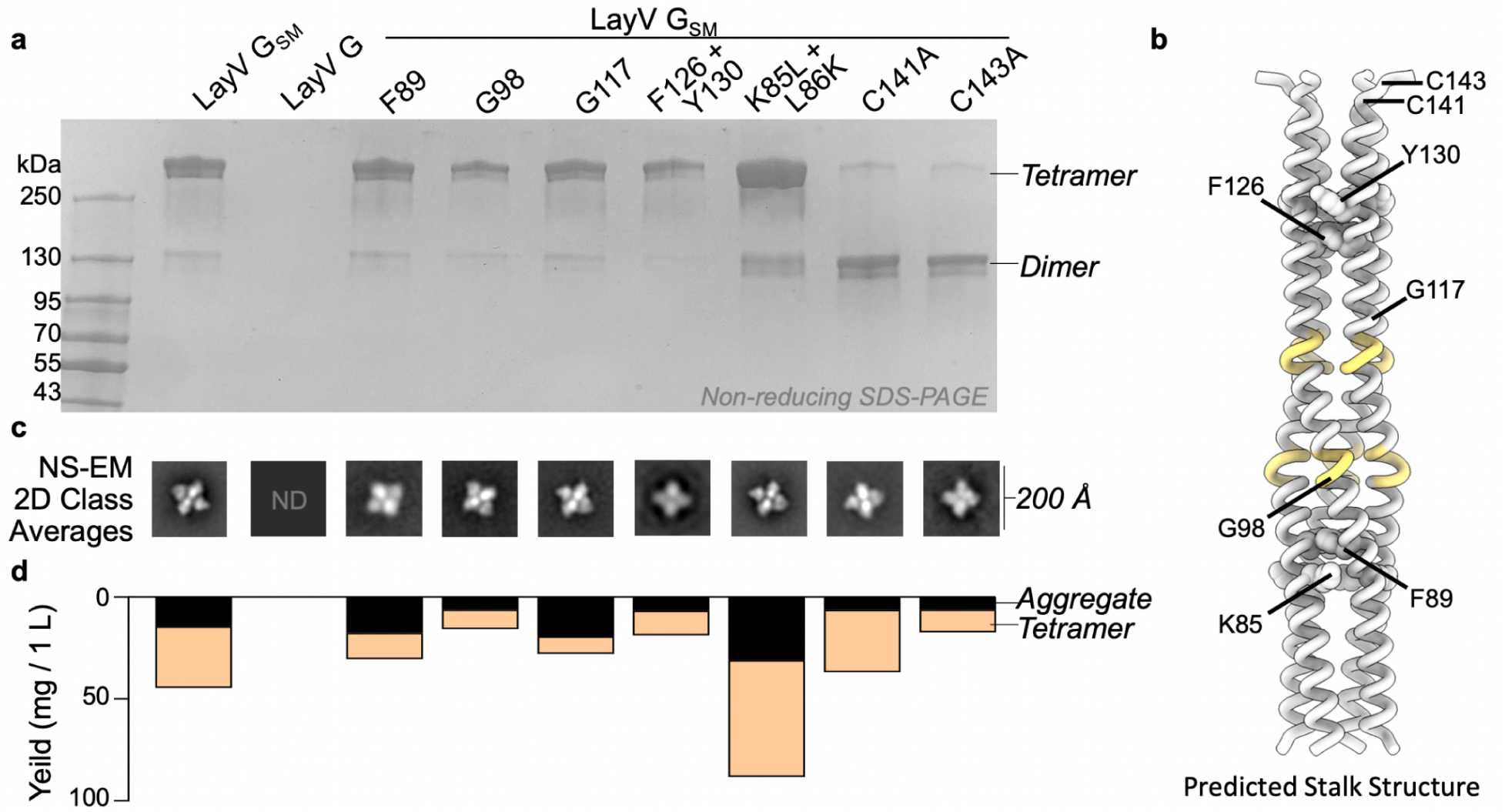
Computational design of a stabilized LayV G tetramer. **a**, Non-reducing SDS-PAGE analysis of purified LayV G mutants after affinity purification, normalized to a standard volume to show the relative purification yield. **b**, Alphafold2 structure prediction of the LayV G tetrameric stalk. Residues with predicted pi-helical secondary structure are shown in yellow. **c,** selected 2D classes that demonstrate each individual mutant is capable of forming a tetramer similar in structure to LayV G_SM_. **d**, Yield of purified LayV G per 1000 mL of cell culture used for protein expression, stratified by tetramers or soluble aggregates as determined by size exclusion chromatography.

To overcome this challenge and improve LayV G expression and stability, we set out to design stabilizing mutations within the G ectodomain (Fig. 5). Tetrameric structure prediction of the LayV G central stalk obtained with AlphaFold2^56^ were overall consistent with the NiV G stalk organization^11^ and suggested that substitutions of three aromatic residues present in the wildtype LayV G stalk (F89, F126, and Y130) could improve core packing (Fig. 5b). Furthermore, two glycine residues (G98 and G117) were anticipated to weaken the local secondary structure of the helical stalk (Fig. 5b). Computational design of optimal residue replacements were aided by ProteinMPNN^57^ and visually inspected to generate a LayV G construct, designated LayV G_SM_, comprising the F89L, G98T, G117A, F126V, and Y130L residue substitutions. Transient transfection of Expi293 human cells with the LayV G_SM_ construct yielded 30 mg of purified protein per liter of culture forming well-folded tetramers with four head domains clustered around a central stalk, as visualized by negative staining EM (Fig. 5 and Extended Data Fig. 8). LayV G_SM_ harboring the L89F, T98G, A117G, or V126F/L130Y reversions had 30-60% reduced yields (Fig. 5d) indicating that the designed mutations act collectively to stabilize the ectodomain tetramer.

### Architecture of LayV G

To unveil the 3D organization of the LayV G glycoprotein tetramer, we determined a 3.2 Å resolution cryoEM structure of LayV G_SM_ (Fig. 6a**-b** and Extended Data Fig. 9 and Table **1**). LayV G folds as a 150 Å-high and 140 Å-wide 4-fold symmetrical tetramer with a central stalk surrounded by four β-propeller head domains (Fig. 6a**-b**). The stalk region built in the cryoEM density comprises residues 72 to 142 and the β-propeller head domain comprises residues 176 to 607. Residues 166 to 175 of the intervening linker region is resolved in the LayV G map and appears to mediate interprotomer interactions with a neighboring head domain (Fig. 6a). The neck region approximately spanning residues 143 to 165 was not visible in the map and could not be modeled. The LayV G map resolves a single N-linked glycan at position N189 whereas the N619 glycan is not resolved in the cryoEM density and the N61 and N64 glycans were not included in the ectodomain construct.

**Figure 6:**
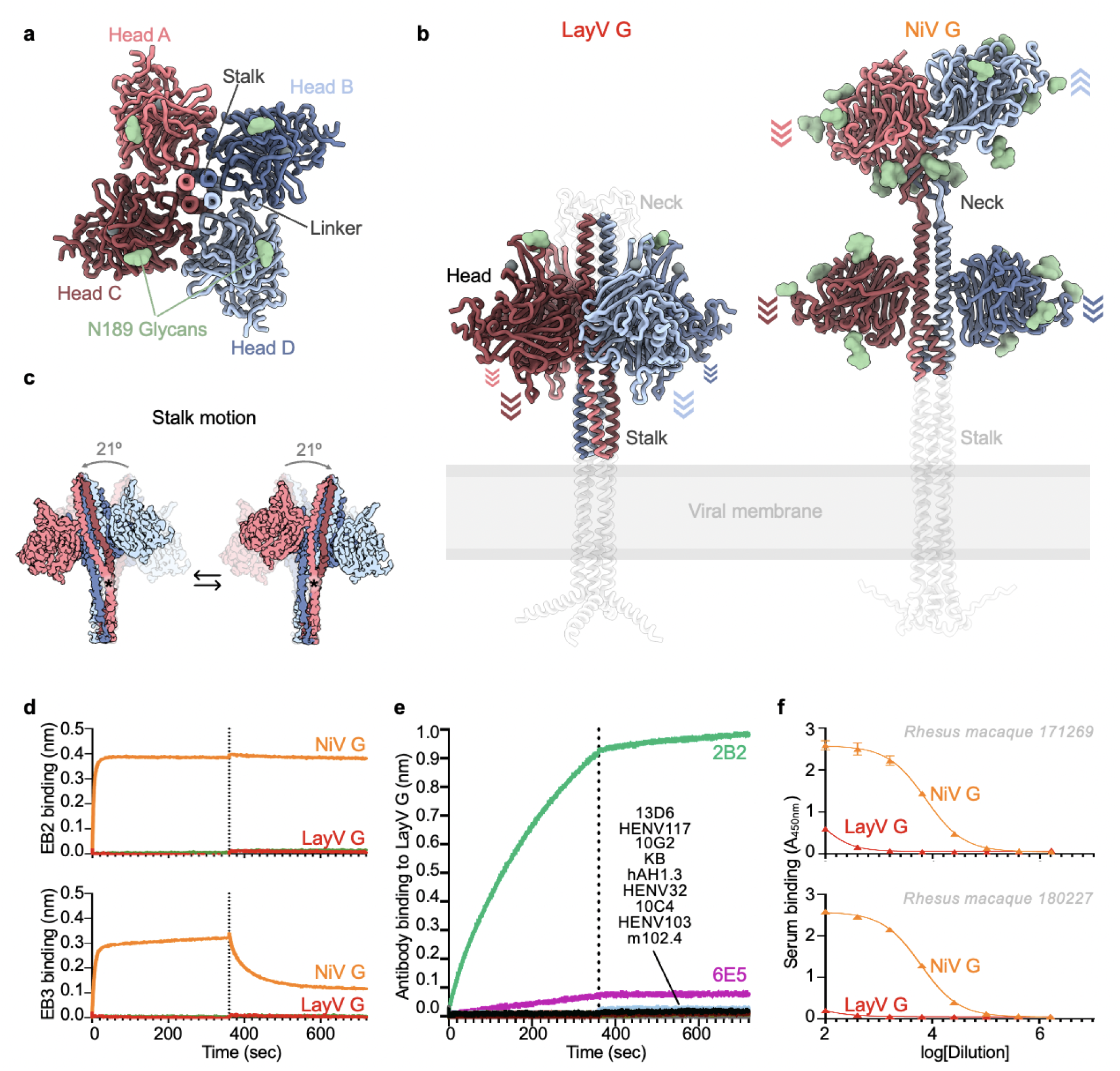
Architecture of the LayV G attachment protein. a, Ribbon diagram of the LayV G ectodomain tetramer shown along the central stalk. **b**, LayV G (left) and NiV G (right, composite from PDB 7TXZ and PDB 7TY0) ectodomain structures shown normal to the central stalk. The Alphafold2 predicted structures of the stalk and neck regions as well as of the transmembrane and cytoplasmic domains that were not empirically determined are shown as transparent white ribbons for reference. Arrows indicate the orientation of each head domain. **c**, Conformational flexibility of LayV G around its central stalk region revealed through cryoSPARC 3DFlex analysis of our cryoEM dataset. Residue K85 is marked with an asterisk. **d,** Binding evaluation of 200nM human EB2-Fc and EB3-Fc to LayV G (red) or NiV G (orange) immobilized at the surface of HIS1K biolayer interferometry biosensors. Negative control (green) is buffer only. **e,** Evaluation of binding of G-directed mAbs at a concentration of 200nM to LayV G or NiV G immobilized at the surface of HIS1K biolayer interferometry biosensors. KB refers to negative control with buffer only (kinetics buffer). **f,** Dose-response curves for binding of rhesus macaque polyclonal serum elicited by equimolar mixture of NiV-M and NiV-B antibodies to LayV G or NiV G using ELISA.

The four LayV G head domains form a 4-fold symmetrical clamp around the central stalk with each β-propeller interacting with two neighboring heads and with the stalk helices from two distinct protomers (Fig. 6a**-b**). The connectivity between the stalk and heads is not resolved, due to the neck disorder, so each head was modeled as the same chain as the stalk chain it primarily interacts with (by analogy with NiV G^11^). The LayV G head domain conformation observed in our structure markedly differs from that found in NiV G as the latter glycoprotein adopts an asymmetric organization of its head domains with only two of them interacting with the stalk (Fig. 6b). The orientation of the four LayV G heads is roughly reminiscent of the two NiV G heads positioned against the stalk, i.e. with the NiV G ephrin B2-binding site pointing towards the viral membrane, although the exact head domain positioning relative to the stalk is different. The observed LayV G head conformation and symmetry is unique among characterized paramyxovirus attachment glycoprotein structures (Extended Data Fig. 10).

Although LayV G_SM_ mainly migrates as a tetramer when analyzed by non-reducing SDS-PAGE, purification of LayV G_SM_ harboring the C141A or the C143A neck residue mutations led to purification of covalent dimers (Fig. 5b). These data indicate that these two cysteine residues separately promote covalent LayV G dimerization and that LayV G is a dimer of dimers, as is the case for NiV G and HeV G ^58^, although the organization of their neck regions is distinct. The stalk region is a four-helix coiled coil with intervening pi-helices at residues 97-101 and 111-115 mediated by prolines at residues 102 and 116, consistent with the NiV G stalk region (Fig. 6b and Extended Data Fig. 11). Twenty additional residues could be resolved at the N-terminal region of the stalk, relative to the NiV G structure, revealing an additional pi-helix followed by a 3_10_-helix at residues 82-89. 3DFlex analysis^59^ of the cryoEM data using cryoSPARC demonstrates this pi-helix and 3_10_-helix combination creates a hinge-like pivot point in the stalk permitting a ∼20° rotation of the membrane-proximal region relative to the head-interacting region of the stalk (Fig. 6c and Extended Data video **1**). Given the observed LayV G stalk flexibility, host-receptor binding could possibly occur on the face of the head domain facing toward or that pointing away from the viral membrane.

Structural analysis supports that K85 and L86 residues mediate this pi-helix and 3_10_-helix combination and can be optimized for improved protein expression. To maintain alpha helical integrity, K85 would need to orient towards and L86 away from the core of the coiled coil, which would position K85 in a hydrophobic environment and L86 in a hydrophilic environment. Instead, we observed that K85 is oriented away from and L86 towards the core of the coiled coil, positioning K85 in a hydrophilic environment and L86 in a hydrophobic environment (Extended Data Fig. 11). This break from alpha-helical structure creates a pi-helix before K85/L86 and a 3_10_-helix after K85/L86 (Extended Data Fig. 11). These residues are conserved in MojV G, GAKV G, and DARV G and would be expected to play a similar role (Extended Data Fig. 11). In distantly related HNVs, K85 and L86 are hydrophobic and hydrophilic residues, respectively (Extended Data Fig. 11). Therefore, we expect that K85L and L86K mutations stabilize local alpha-helical secondary structure and addition of these mutations to LayV G_SM_ enhances the production yield 2-fold (Fig. 5d).

### LayV G is functionally and antigenically distinct from NiV G

Due to the close evolutionary relationship between LayV and MojV G glycoproteins along with their marked divergence from NiV G and HeV G, we postulated that LayV G would not engage ephrin B2 or B3 (NiV/HeV host receptors^24,26^). Indeed, we could not detect binding of human ephrin B2 or B3 to LayV G using biolayer interferometry (Fig. 6d). Furthermore, a panel of NiV/HeV G-directed neutralizing mAbs (m102.4, HENV-32, HENV-103, HENV-117, nAH1.3), a NiV G-specific mAb (13D6) and a HeV G-specific mAb (hAH1.3) also failed to recognize LayV G (Fig. 6e). However, we found that out of three MojV G-elicited mAbs (6E5, 10G2, 2B2), one of them (2B2) strongly cross-reacted with LayV G and another one (6E5) exhibited weak binding (Fig. 6e). These data suggest that LayV G has a different receptor tropism from previously characterized HNVs (as is the case for MojV) and distinct antigenicity, as the mAb cross-reactivity analysis mirrors that carried out with LayV F-directed antibodies.

Based on these results along with the large genetic distance between LayV G and NiV/HeV G and the conformational differences observed between them, we hypothesized that vaccine-elicited antibodies based on NiV G or HeV G might not be cross-reactive with LayV G. To explore this possibility, we used sera from two rhesus macaques that were immunized with an equimolar mixture of purified NiV-Bangladesh and NiV-Malaysia G ectodomain tetramers^11^. Using ELISA, we could not detect appreciable antibody binding to LayV G with any of the two non-human primate sera at a dilution of 1/100 whereas we observed strong antibody binding titers against NiV G (Fig. 6f). These findings support and extend the lack of cross-reactivity of NiV/HeV G-elicited mAbs and indicate that vaccine candidates currently being developed against NiV and HeV would not elicit neutralizing antibodies against LayV G, which share only 22% amino acid sequence identity with NiV G.

## Discussion

Our structures of prefusion LayV F and LayV G reveal the ultrastructural organization of these key targets of the immune system and provide blueprints explaining their distinct antigenicity relative to NiV and HeV at the molecular level, as illustrated by the lack of cross-reactivity of NiV/HeV-elicited mAbs. The LayV G conformation observed in our cryoEM structure, with all four head domains clamped down along the stalk, and the intrinsic structural flexibility detected have not been previously described for any paramyxovirus attachment glycoproteins and may represent snapshots of the cascade of conformational changes leading to activation of F for membrane fusion and initiation of infection. Our postfusion F structure represents the ground state of the fusion reaction, as illustrated by the large enhancement of buried surface area at the interface between protomers relative to prefusion F. Furthermore, postfusion F not only resembles other paramyxovirus F trimers but also postfusion coronavirus S trimers, thereby adding to the body of data supporting a common evolutionary origin and mechanism of action for the fusion machinery of these two viral families^41^.

The cross-neutralizing NiV and HeV G-specific m102.4 mAb has been administered on an emergency compassionate use basis to 15 individuals with high-risk exposure to HeV or NiV and recently completed a phase 1 clinical trial in Australia. m102.4 is the sole antibody evaluated in the clinic against any HNV^60^ and we showed here that it failed to bind to LayV G due its low sequence conservation with NiV G (22% amino acid identity). Moreover, a vaccine based on the HeV G tetrameric ectodomain has been shown to protect against both NiV and HeV challenge in preclinical studies and is commercially available for use in horses in Australia (Equivac HeV, Zoetis Inc.) ^21,23,61–65^. Although a similar immunogen adjuvanted with alum has recently entered phase 1 clinical trials (NCT04199169), we demonstrate here that NiV G vaccine-elicited antibodies, that are neutralizing NiV, HeV and the recently emerged divergent HeV-g2^11,52^, did not appreciably cross-react with LayV G. Collectively, our data strongly indicate that countermeasures being advanced to the clinic will be ineffective against LayV G which has already been detected in several individuals and therefore represent a current threat to public health^31^. Our data urge for the development of vaccines and mAb therapies against LayV G and closely related HNVs (e.g MojV, DARV and GAKV).

Structure-guided viral glycoprotein engineering has revolutionized vaccine design against a range of emerging or endemic pathogens. Stabilization of the respiratory syncytial virus fusion glycoprotein in its prefusion conformation (DS-Cav1) propelled the development of a safe and effective vaccine against this pathogen^66–68^. Prefusion-stabilization (2P) of coronavirus spike glycoproteins also led to the groundbreaking development of COVID-19 vaccines in a record time in 2020^49,69–71^. We describe here the computational design of LayV G constructs, enabling high-yield recombinant production of the LayV G ectodomain tetramer, and show that a disulfide stapling HNV F trimers in the prefusion state^13,17^ is a broadly applicable prefusion-stabilization strategy yielding monodisperse LayV F and GhV F trimers. These protein design strategies will enable the preclinical evaluation of next-generation vaccine candidates against the newly emerged LayV.

## METHODS

### Cell lines

Expi293F is a derivative of the HEK293 cell line (Thermo Fisher). Expi293F cells were grown in Expi293 Expression Medium (Thermo Fisher), cultured at 37°C with 8% CO_2_ and 130 rpm. Vero 76 (ATCC CRL-1587) cells were maintained at 37°C, 5% CO_2_ in Dulbecco’s modified eagle media (DMEM) (Quality Biological; MD, USA) supplemented with 10% cosmic calf serum (CCS) and 1% L-glutamine (Quality Biological; MD, USA) (DMEM-10). CHO-K1 cells (ATCC CCL-61) were maintained in Ham’s F-12K (Kaighn’s) Medium (Gibco) with 1% penicillin–streptomycin and 10% CCS at 37°C with 5% CO_2_. HEK293T cells and Neuro-2a cells (ATCC CCL-131) were maintained in DMEM supplemented with 2 mM L-glutamine, 1% penicillin–streptomycin (Quality Biologicals) and 10% CCS. HEK293T cells were maintained at 37°C with 5% CO_2_, while Neuro-2a cells were maintained at 37°C with 8% CO_2_.

### Recombinant LayV and GhV F ectodomain production

The LayV F ectodomain construct used for all experiments in this study includes codon optimized LayV F (residue 1-482) fused to a C-terminal GCN4 followed by a factor Xa sequence, a linker (GSGGGS) and an hexa-histidine tag (HHHHHH) into pTwist CMV vector for transient expression using Expi293F cell line. The GhV F ectodomain construct used for all experiments in this study includes codon optimized GhV F used for structure determination comprise residue 1-586 and is C-terminally fused to the I53-50A component via an intervening 16 GS linker (as previously described^72^) followed by a GSGGGS linker and a 6×His tag. GhV F wildtype is composed of residue 1-586 with a Cterminal GSGGGS linker followed by a 6×His tag. The LayV F wildtype, LayV F N95C/A114C, LayV F I167F/S186P, LayV F I167F/S186P/N95C/A114C, GhV F wildtype, GhV F S196C/A215C, GhV F I268F/Q287P, GhV F I268F/Q287P/S196C/A215C and GhV F I50-50A, were synthesized by Twist Bioscience. LayV F and GhV F ectodomains were produced in 100mL Expi293F cells grown in suspension using Expi293 Expression Medium (Thermo Fisher) at 37°C in a humidified 8% CO_2_ incubator rotating at 130 rpm. The cultures were transfected using ExpiFectamine™ 293 Transfection Kit (Thermo Fisher) with cells grown to a density of 3 million cells per mL and cultivated for 4 days. The supernatants were harvested, and proteins were purified from clarified supernatants using a 1mL HisTrap HP column (Cytiva), buffer exchanged, concentrated and flash frozen in either TBS (50 mM Tris, 150 mM NaCl and 10 mM EDTA at pH 8.0). SDS-PAGE was run to check purity.

### Recombinant LayV G ectodomain production

LayV G ectodomains were produced in 25 mL culture of Expi293F Cells (ThermoFisher Scientific) grown in suspension using Expi293 Expression Medium (ThermoFisher Scientific) at 37°C in a humidified 8% CO_2_ incubator rotating at 130 rpm). Cells grown to a density of 3 million cells per mL were transfected with the ExpiFectamine 293 Transfection Kit (ThermoFisher Scientific) and cultivated for four days at which point the supernatant was harvested. LayV G was purified from clarified supernatants using 2 mL of cobalt resin (Takara Bio TALON), washing with 200 column volumes of 50 mM Tris-HCl pH 8.0, 150 mM NaCl, and 5 mM imidazole, and eluted with Tris-HCl pH 8.0, 150 mM NaCl, and 600 mM imidazole. Eluted protein was further purified by size exclusion chromatography using a Superose 6 increase 10/300 GL column (Cytiva) equilibrated in a buffer containing Tris-HCl pH 8.0 and 150 mM NaCl. Purified protein was concentrated using a 100 kDa centrifugal filter (Amicon Ultra 0.5 mL centrifugal filters, MilliporeSigma) to 2 mg/mL and flash frozen with liquid nitrogen. The LayV G_SM_ ectodomain has a human azurocidin signal peptide (MTRLTVLALLAGLLASSRA) followed by an N-terminal His-tag (SGPHHHHHHHHGSSP) followed by residues 72 through 624 of LayV G. The LayV G_SM_ ectodomain also has F89L, G98T, G117A, F126V, and Y130L mutations, unless otherwise denoted in the text. The LayV G_SM_ ectodomain used for structure determination also had a designed tetramerization sequence (QAEELKAIKEELKAIKEELKAIKEELKAI) between the His-tag and residue 72 of LayV G for further stabilization; though this was subsequently found to be unnecessary as LayV G_SM_ is stable on its own.

### MojV F and G-reactive antibody generation

Codon-optimized MojV (Genbank NC 025352.1) F and G open reading frames (ORFs) were synthesized by GenScript® (Piscataway, NJ, USA) and subcloned into a pcDNA3.1 Hygro CMV mammalian expression vector. For construction of soluble and secreted versions of F glycoproteins, the transmembrane domain (486-520) and C-terminal cytoplasmic tail (521-545) of MojV F were replaced with a GCN4 trimerization motif (GCNt) (MKQIEDKIEEILSKIYHIENEIARIKKLIGE), followed by a factor Xa protease cleavage site (IEGR) and an S-tag to generate trimeric soluble construct of MojV sF. To produce tetrameric soluble MojV G (MojV sG), the N-terminal cytoplasmic tail (2-37) and transmembrane domain (38-70) were replaced with an Ig*k* leader sequence (METDTLLLWVLLLWVPGSTGD) and an S-tag followed by a factor Xa protease cleavage site (IEGR) and a GCN4 tetramerization motif (GCNtet) (MKQIEDKLEEIESKLKKIENELARIKK).

For protein expression and purification,human FreeStyle™ 293F cells were transfected with sF, and Neuro-2a cells with sG plasmid constructs. Their stable cell lines were obtained through two rounds of limiting dilutions. Stable expressed MojV sF and sG secreted into FreeStyle™ 293 Expression Medium were collected and firstly purified through a S-agarose affinity column (XK 26 column). For MojV sG, the column was washed with 3x bed volumes of Buffer I (PBS, 0.1% Triton® X-100, 0.3 M NaCl), followed by 6x bed volumes of Wash Buffer II (PBS, 0.1% Triton® X-100). The protein was eluted with Elution buffer (0.2 M citrate, pH 2), then neutralized with buffer (1 M Tris-HCl pH 8.0 or 1 M HEPES buffer pH 9.0). The protein was buffer exchanged to PBS, pH7.4. For MojV sF, the column was washed with 3x bed volumes of Wash buffer I (PBS, 0.5% Triton X-100, 0.5 M NaCl, 0.1 M L-Arginine, 0.02 M Tris-HCl, pH 7.5. followed by 6x bed volumes of Wash buffer II (PBS, 0.1% Triton® X-100, 0.1 M L-Arginine). The protein was eluted with Elution buffer (0.2 M citrate, 0.2 M L-Arginine, pH 2) and neutralized with buffer (1 M HEPES buffer pH 9.0). The protein was buffer exchanged into PBS, 0.01% Triton® X-100, 0.1 M L-Arginine. S-tag was removed with Factor Xa Cleavage Capture kit (Millipore Sigma). The untagged sF or sG was further purified with size-exclusion column (HiLoad 16/60 Superdex 200 prep grade XK 16 gel filtration column (GE Healthcare). The buffer PBS was used for sG and the PBS with 0.01% Triton X-100 was used for sF purification.

To generate F-or G-directed monoclonal antibodies, purified sF_GCNt_ or sG_GCNtet_ glycoproteins were used to immunize BALB/c mice purchased from Jackson Labs (Bar Harbor, ME, USA). All animal studies were carried out under an approved protocol MIC-16-262 for animal experiments obtained from the Uniformed Services University Animal Care and Use Committee. All mice were immunized 4 times.The administered antigens (sF/sG) were formulated in a Sigma adjuvant system (Sigma-Aldrich Co. LLC, St. Louis, MO). Mouse splenic lymphocytes were isolated 4 days following a final immunization without adjuvant and fused with SP2/0 cells using polyethylene glycol by standard methods. Hybridoma supernatants producing MojV F (4G5) or G(2B2)-specific antibodies were identified by enzyme-linked immunosorbent assay (ELISA) using purified S-peptide-cleaved MojV sF or sG. The anti-F or G mAbs were prepared under serum-free conditions using HyClone SFM4MAb (Thermo Fisher Scientific Inc., Rockford, IL) and purified by protein G-Sepharose affinity chromatography.

### Antibody production

nAH1.3 heavy chain and light chain were codon optimized and synthesized by GeneArt (2019AAG4YC) with a signal sequence (MPMGSLQPLATLYLLGMLVASVLA) at the N-termini of both heavy and light chains, as well as a StrepII tag (WSHPQFEK) linked by a linker sequence (GSGGGS) at the C terminal of the heavy chain. The synthesized heavy and light chain genes were each subcloned into a pcDNA3_1(+)_MAr vector. nAH1.3 IgG was produced in 25mL Expi293F cells grown in suspension using Expi293 Expression Medium (Thermo Fisher) at 37°C in a humidified 8% CO2 incubator rotating at 130 rpm. The cultures were double transfected with 1:1 ratio of heavy chain and light chain plasmids using ExpiFectamine™ 293 Transfection Kit with cells grown to a density of 3 million cells per mL and cultivated for 6 days. The supernatants were harvested, and proteins were purified from clarified supernatants using a 1mL StrepTrap™ HP column (Sigma), buffer exchanged, concentrated and flash frozen in either TBS or PBS. SDS-PAGE was run to check purity.

HENV-32, HENV-103, and HENV-117 IgG genes were codon-optimized for a mammalian cell expression system, synthesized, and cloned into a pCDNA3.1+ vector by GenScript. The HENV-32 light-chain sequence was obtained from the Fab sequence of crystal structure Protein Data Bank (PDB 6VY4, and the HENV-32 heavy-chain sequence was reconstructed by combining the Fab sequence of crystal structure PDB 6VY4 with a consensus sequence of IGHG1 (P01857). nAH1.3 and HENV-32 heavy chains include a StrepII tag (WSHPQFEK) linked by a linker sequence (GSGGGS) at the C terminus. HENV-103 and HENV-117 light-and heavy-chain constructs have been produced by replacing the variable domain of HENV-32 constructs with the corresponding HENV-103 or HENV-117 sequence. The C-terminal StrepII tag and linker have been removed for HENV-103 and HENV-117 heavy-chain constructs.

m102.4 was isolated from a recombinant human phage-displayed Fab library and prepared as previously described^55^. Briefly, CHO-K1 cells were transfected with a linearized m102.4 PDR12 construct and the antibody was purified with protein A from a selected stable m102.4-producing cell line^55^. The human IgG1 m102.4 mAb has been extensively characterized as a highly potent HeV and NiV cross-reactive _mAb_^2^2,54,55,60,73,7^4^.

5C4, 5B3, 12B2, 6E5, 13D6, 10G2, hAH1.3 IgGs were produced from hybridomas and were previously described as^36,46^.

### Biolayer interferometry

Assays were performed on an Octet Red (ForteBio) instrument at 30°C with shaking at 1,000 RPM.

For anti HNV mAbs screening, Anti-penta His (HIS1K) biosensors were hydrated in water for 10 min prior to a 120 s incubation in 10X kinetics buffer (undiluted). LayV F N95C/A114C or LayV G central stalk stabilized ectodomain was loaded at 10 μg/mL in 10X Kinetics Buffer for 360 s prior to baseline equilibration for 300 s in 10X kinetics buffer. Association of anti HNV IgGs in 10X kinetics buffer at 200nM concentration was carried out for 360 s prior to dissociation for 360 s.

For 4G5 K_D_ determination of prefusion LayV F, anti-penta His (HIS1K) biosensors were hydrated in water for 10 min prior to a 120 s incubation in 10X kinetics buffer (undiluted). LayV F N95C/A114C was loaded at 20 μg/mL in 10X Kinetics Buffer for 300 s prior to baseline equilibration for 300 s in 10X kinetics buffer. Association of 4G5 Fab in 10X kinetics buffer at various concentrations in a two-fold dilution series from 100nM to 3.125nM was carried out for 100 s prior to dissociation for 100 s.

For 4G5 K_D_ determination of postfusion LayV F (spontaneous refolded over 4 months at 4°C), anti-penta His (HIS1K) biosensors were hydrated in water for 10 min prior to a 120 s incubation in 10X kinetics buffer (undiluted). LayV F spontaneous refolded was loaded at 20 μg/mL in 10X Kinetics Buffer for 300 s prior to baseline equilibration for 300 s in 10X kinetics buffer. Association of 4G5 Fab in 10X kinetics buffer at various concentrations in a two-fold dilution series from 40nM to 1.25nM was carried out for 360 s prior to dissociation for 360 s.

For 4G5 K_D_ determination, the data were baseline subtracted prior to fitting performed using a 1:1 binding model and the ForteBio data analysis software. Mean kon, koff values were determined with a global fit applied to all data. The experiments were done with two separate purification batches. Data was plotted and fit in Prism (GraphPad).

### Enzyme-linked immunosorbent assay (ELISA)

96-well Maxisorp plates (Thermo Fisher) were coated overnight at 4°C with 2 µg/mL of NiV G or LayV G central stalk stabilized ectodomain in 50mM Tris and 150mM NaCl at pH 8. Plates were slapped dry, washed 3X in Tris Buffered Saline Tween (TBST) and blocked with Blocker Casein (ThermoFisher) for 1 h at 37°C. Plates were slapped dry and washed 4X in TBST. 1:4 serial dilutions of NHP sera were made in 50 μL TBST and incubated at 37°C for 1 h. Plates were slapped dry and washed 4X in TBST followed by addition of 50 μL 1:5000 Goat anti-Human IgG Fc Secondary Antibody with HRP (Invitrogen) for one hour at 37°C. Plates were slapped dry and washed 4X in TBST followed by addition of 50 μL TMB Microwell Peroxidase (Seracare). The reaction was quenched after 4 minutes with 1 N HCl and the A450 of each well was read using a BioTek Synergy Neo2 plate reader. Data was plotted and fit in Prism (GraphPad) using nonlinear regression sigmoidal, 4PL, X is log(concentration).

### Negative stain EM

3 μl of purified sample of each F constructs was applied to a glow discharged 300 mesh copper grids (Ted Pella) coved with evaporated continuous carbon on top of collodion (Electron Microscope Science) layer for 1 min before blotting away excess liquid with Whatman no. 1 filter paper. Grids were stained with 3 μl of 2% (w/v) uranyl formate stain for 1 min twice. Grids were imaged using an FEI Tecnai Spirit 120 kV electron microscope equipped with a Gatan Ultrascan 4000 CCD camera. The pixel size at the specimen level was 1.60 Å. Data collection was performed using Leginon^75^. All data were processed using CryoSPARC^76^ following the pipeline described in Extended Data Fig. 1.

### LayV F and G expression plasmids and Peptide synthesis

Codon optimized LayV F and G open reading frames were synthesized by Genscript and subcloned into a mammalian promoter modified expression vector pcDNA3.1-CMV^77^. To enable detection, an S peptide tag (KETAAAKFERQHMDS) was inserted at the C-terminal end of LayV F and at the N-terminal end of LayV G. A peptide corresponding to the heptad repeat 2 (HR2) domain of LayV F (LayV HR2, KIDIGNQLAGINQTLQNAEDYIEKSEEFLKGINPSI) and the corresponding scrambled peptide (scrambled LayV HR2, SIANIQEKDIIKLETEDPEIYAGNKLGSQILNFGQN) were synthesized by Biosynth (Gardner, MA, USA).

### Western blot

CHO-K1, HEK293T and Neuro-2a cells were transfected with pcDNA3.1-LayV F, pcDNA3.1-LayV G, pcDNA3.1-LayV F and pcDNA3.1-LayV G encoding S-tagged constructs. As a reference, cells were transfected with pcDNA3.1-MojV F, pcDNA3.1-MojV G or pcDNA3.1-MojV F and pcDNA3.1-MojV G encoding S-tagged constructs^36^. At 48 hrs post transfection, cells were lysed with 1x RIPA (radioimmunoprecipitation assay) Lysis and Extraction Buffer containing a protein inhibitor cocktail and incubated with S protein agarose beads at 4°C overnight. Proteins were resolved by SDS-PAGE and detected with an anti-S-peptide-HRP conjugated antibody.

### Cell-cell fusion assay

We used a quantitative dual-split-reporter luciferase-based cell-cell fusion assay as previously described^78^. Briefly, the effector cells (Neuro-2a cells) were transfected with pcDNA3.1-LayV F, or pcDNA3.1-LayV G or pcDNA3.1-LayV F and pcDNA3.1-LayV G together with one half of the dual-split-reporter luciferase expression plasmid (DSP1–7). As a control, Neuro-2a cells were co-transfected with pcDNA3.1-MojV F, pcDNA3.1-MojV G and DSP 1-7. Each target cell (HEK293T or Neuro-2a cells, endogenously expressing HNV receptors) were transfected with the other half of the dual-split reporter plasmid (DSP 8–11). After 48 hours post-transfection, the effector cells were gently detached and applied onto the target cells in equal numbers. At 48 hrs post mixing, EnduRen (Live cell substrate for the luciferase) was added to the plate. Content mixing between the effector cells and the target cells as a result of cell-cell fusion was measured using a luminometer (as RLU, Relative Luminescence Unit). Each assay was done for three replicates. Cell-cell fusion inhibition was assessed by LayV HR2 and scrambled LayV HR2 peptides.

### CryoEM sample preparation and data collection

To prepare the CryoEM grids of LayV F for solving prefusion state, 3 μL of the 0.14 mg/mL LayV F wildtype were loaded onto holey carbon grids covered with graphene oxide, prior to plunge freezing using a Vitrobot MarkIV (ThermoFisher Scientific) with a blot force of -1 and 3 second blot time at 100 % humidity and 22°C. Data were acquired using an FEI Titan Krios transmission electron microscope operated at 300 kV and equipped with a Gatan K3 direct detector and Gatan Quantum GIF energy filter, operated in zero-loss mode with a slit width of 20 eV. The dose rate was adjusted to 15 counts/pixel/s, and each movie was fractionated in 75 frames of 40 ms. Automated data collection was carried out using Leginon at a nominal magnification of 105,000x with a pixel size of 0.843 Å in defocus range between -0.7 and -1.7 μm^75^.

To prepare the CryoEM grids of LayV F for solving postfusion state, 3 μL of 2.7 mg/ml LayV F wildtype (spontaneous refolded to postfusion over 3 months at 4°C) with 10mM OG detergent were applied onto a freshly glow discharged 2.0/2.0 UltraFoil^79^ grid (200 mesh), plunge frozen using a vitrobot MarkIV (ThermoFisher Scientific) using a blot force of 0 and 6 second blot time at 100 % humidity and 22°C. Data were acquired using the SerialEM to control a Glacios transmission electron microscope equipped with a Gatan K3 Summit direct detector and operated at 200 kV. The dose rate was adjusted to 7.5 counts/pixel/s, and each movie was acquired in 100 frames of 50 ms with a defocus range between −0.7 and −1.7 μm and a pixel size of 0.89 Å^80,81^.

To prepare the CryoEM grids of LayV F in complex with 4G5 Fab, 3 μL of 1 mg/ml LayV F wildtype (spontaneous refolded to postfusion over 4 months at 4°C) with 10mM OG detergent were applied onto a freshly glow discharged 2.0/2.0 UltraFoil^79^ grid (200 mesh), plunge frozen using a vitrobot MarkIV (ThermoFisher Scientific) using a blot force of 0 and 6 second blot time at 100 % humidity and 22°C. Data were acquired using an FEI Titan Krios transmission electron microscope operated at 300 kV and equipped with a Gatan K3 direct detector and Gatan Quantum GIF energy filter, operated in zero-loss mode with a slit width of 20 eV. The dose rate was adjusted to 15 counts/pixel/s, and each movie was fractionated in 75 frames of 40 ms. Automated data collection was carried out using SerialEM at a nominal magnification of 105,000x with a pixel size of 0.843 Å in defocus range between -1.3 and -1.7 μm^80,81^.

To prepare the CryoEM grids of LayV G, 3 μL of 0.6 mg/mL LayV G or 2 mg/ml LayV G with 0.01% fluorinated octyl-maltoside (Anatrace) detergent were applied onto a freshly glow discharged 2.0/2.0 UltraFoil^79^ grid (200 mesh), plunge frozen using a vitrobot MarkIV (ThermoFisher Scientific) using a blot force of −1 and 6 second blot time at 100% humidity and 23°C. Data were acquired using the Leginon software^75^ to control a Glacios transmission electron microscope equipped with a Gatan K3 Summit direct detector and operated at 200 kV. The dose rate was adjusted to 7.5 counts/pixel/s, and each movie was acquired in 100 frames of 50 ms. For LayV G with and without Fluorinated Octyl Maltoside detergent, 4894 and 4086 micrographs, respectively, were collected in a single session with a defocus range between −0.5 and −2.4 μm and a pixel size of 0.89 Å.

To prepare the CryoEM grids of GhV F I53-50A, 3 μL of the 0.05 mg/mL GhV F I53-50A were loaded onto holey carbon grids covered with graphene oxide, prior to plunge freezing using a Vitrobot MarkIV (ThermoFisher Scientific) with a blot force of -1 and 3 second blot time at 100 % humidity and 22°C. For the data set of GhV F, data were acquired using an FEI Titan Krios transmission electron microscope operated at 300 kV and equipped with a Gatan K3 direct detector and Gatan Quantum GIF energy filter, operated in zero-loss mode with a slit width of 20 eV. The dose rate was adjusted to 15 counts/pixel/s, and each movie was fractionated in 75 frames of 40 ms. Automated data collection was carried out using Leginon at a nominal magnification of 105,000x with a pixel size of 0.843 Å in defocus range between -0.5 and -2.5 μm^75^.

### CryoEM data processing

For structure of LayV F prefusion, movie frame alignment, estimation of the microscope contrast-transfer function parameters, particle picking, and extraction were carried out using CryoSPARC^76^. Two rounds of reference-free 2D classification were performed using CryoSPARC^76^. 3D refinements were carried out using non-uniform refinement^82^ with reference map from previously solved HNV F.

For structure of LayV F postfusion, movie frame alignment, estimation of the microscope contrast-transfer function parameters, particle picking, and extraction with 2X binning were carried out using CryoSPARC^76^. Reference-free 2D classification was performed using CryoSPARC^76^. Ab initio structure reconstruction was performed in cryoSPARC using three classes^76^. Subset of good postfusion LayV F particles were used for homologous refinement followed by non-uniform refinement. Particles were extracted with original pixel size followed by another round of non-uniform refinement.

For structure of LayV F postfusion in complex with 4G5 Fab, movie frame alignment, estimation of the microscope contrast-transfer function parameters, particle picking, and extraction were carried out using Warp^83^. Two rounds of reference-free 2D classification were performed using CryoSPARC^76^. Heterogeneous refinement was used to sort good particles followed by homologous refinement and non-uniform refinement with per-particle defocus refinement in CryoSPARC^82^. Selected particle images were subjected to the Bayesian polishing procedure^84^ implemented in Relion before performing another round of homologous refinement and non-uniform refinement in cryoSPARC. Subsequently, local non-uniform refinements were performed for the proximal and distal parts of the complex yielding reconstructions at 3.6Å and 3.3Å resolution, respectively.

For the structure of LayV G, movie frame alignment and binning to 1.78 Å was carried out using Warp^83^, estimation of the microscope contrast-transfer function parameters, particle picking, and extraction (with a box size of 440 pixels^2^) was carried out in cryoSPARC^76^. Reference-free 2D classification was performed using cryoSPARC to select well-defined particle images. Ab initio structure reconstruction and non-uniform refinement in cryoSPARC were then performed. 3D classification with 50 iterations each (angular sampling 7.5° for 25 iterations and 1.8° with local search for 25 iterations) were carried out using Relion^85^ without imposing symmetry to select well-defined particle classes. Particle images were then subjected to Bayesian polishing using Relion^84^ during which the box size was adjusted to 440 Å with a pixel size of 0.89 Å. Another round of non-uniform refinement in cryoSPARC was performed concomitantly with global and per-particle defocus refinement as well as beam tilt refinement^82,85^. The particles from this refinement were used with 3DFlex^59^ in cryoSPARC to determine the motions of LayV G, using the default setting for Flex Data Prep, Flex Mesh Prep, Flex Train, and Flex Generate, with the exception that the training box size was set to 220 pixels.

For the structure of GhV F, movie frame alignment, estimation of the microscope contrast-transfer function parameters, particle picking, and extraction were carried out using Warp^83^. Two rounds of reference-free 2D classification were performed using CryoSPARC^76^ to select well-defined particle images. 3D refinements were carried out using non-uniform refinement^82^ along with per-particle defocus refinement in CryoSPARC. Selected particle images were subjected to the Bayesian polishing procedure^84^ implemented in Relion before performing another round of non-uniform refinement in cryoSPARC followed by per-particle defocus refinement. Local resolution estimation, filtering, and sharpening were carried out using CryoSPARC.

Reported resolutions are based on the gold-standard Fourier shell correlation (FSC) of 0.143 criterion and Fourier shell correlation curves were corrected for the effects of soft masking by high-resolution noise substitution^86,87^.

### Cryo-EM model building and analysis

Initial models of both prefusion and postfusion LayV F were built automatically using ModelAngelo^88^. UCSF ChimeraX^89^ and Coot^90,91^ were used for docking and manually building into the cryoEM maps. All models were refined into the cryoEM maps using Rosetta^92–94^. The LayV G model was further refined with Phenix^95^ and ISOLDE^96^ Validation used MolProbity^97^, EMringer^98^, Phenix^95^ and Privateer^99^. Figures were generated using UCSF ChimeraX^89^ and UCSF Chimera^100^.

## Acknowledgements

This study was supported by the National Institute of Allergy and Infectious Diseases (DP1AI158186 and 75N93022C00036 to D.V., and U19AI142764 to C.C.B. and D.V.), a Pew Biomedical Scholars Award (D.V.), an Investigators in the Pathogenesis of Infectious Disease Awards from the Burroughs Wellcome Fund (D.V.), the University of Washington Arnold and Mabel Beckman cryoEM center and the National Institute of Health grant S10OD032290 (to D.V.). D.V. is an Investigator of the Howard Hughes Medical Institute and the Hans Neurath Endowed Chair in Biochemistry at the University of Washington. We thank Yifan Cheng and David Agard for providing the graphene oxide used to make grids.

## Author Contributions

Z.W., M.M., L.Y., M.A., C.B. and D.V. designed the experiments; L.Y., W.S. M.M., and H.V.D., recombinantly expressed and purified glycoproteins. A.P. participated in the early negative staining studies of LayV F. L.Y. performed the fusion assays. Z.W. and D.V. ported F mutations to LayV F and GhV F. Z.W. carried out cryoEM specimen preparation, data collection and processing for all LayV F structures. M.M. designed stabilized LayV G constructs. M.M. carried out cryoEM specimen preparation, data collection and processing of LayV G. H.V.D. designed the GhV F construct used for cryoEM studies. Y.J.P. carried out cryoEM specimen preparation, data collection and processing of GhV F. Z.W., M.M., Y.J.P. and D.V built and refined atomic models. Z.W. performed binding assays. Z.W., M.M and D.V. analyzed the data and wrote the manuscript with input from all authors; C.B., and D.V. supervised the project.

## Competing Interests

M.M. and D.V. are inventors on patent applications submitted by the University of Washington related to LayV G stabilization. The remaining authors declare that the research was conducted in the absence of any commercial or financial relationships that could be construed as a potential conflict of interest.

---

**Extended Data Fig. 1.**
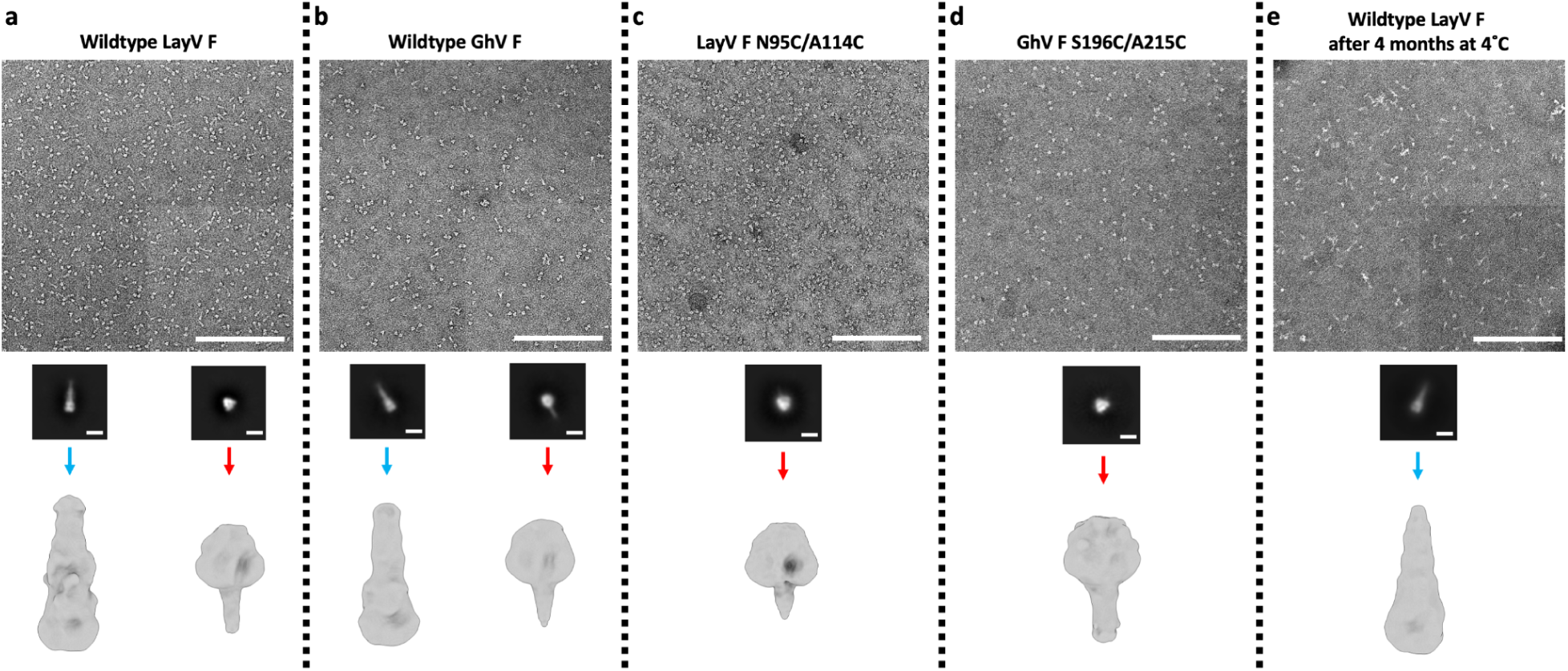
EM data processing pipeline of negatively stained F glycoproteins. **a-e,** Negative stain EM processing pipeline for wildtype LayV F (a), wildtype GhV F (b), LayV F N95C/A114C (c), GhV F S196C/A215C (d) and wildtype LayV F spontaneous refolded to the postfusion state (e). Representative micrographs and 2D class averages are shown. The scale bar represents 200 nm for micrographs or 100 Å for 2D classes. Homo Refine: cryoSPARC homogeneous refinement.

**Extended Data Fig. 2.**
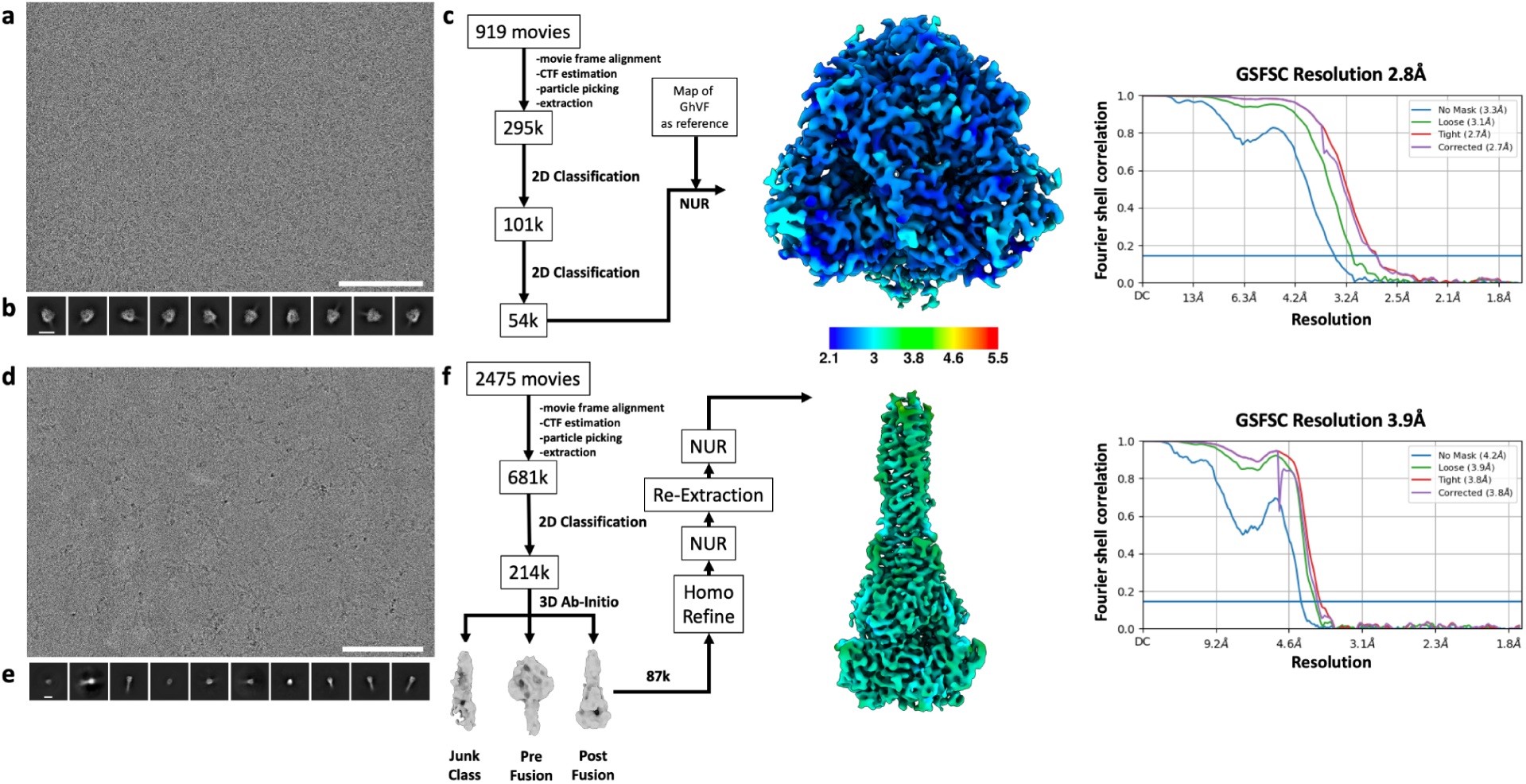
CryoEM data processing pipeline for wildtype prefusion and postfusion LayV F. **a-b,** Representative electron micrograph (a) and 2D class averages (b) of prefusion LayV F embedded in vitreous ice. The scale bar represents 100 nm (a) or 100 Å (b). **c,** CryoEM data processing flow chart for prefusion LayV F, including local resolution maps computed using cryoSPARC. **d-e,** Representative electron micrograph (d) and 2D class averages (e) of postfusion LayV F. The scale bar represents 100 nm (a) or 100 Å (b). **f,** CryoEM data processing flow chart for postfusion LayV F, including local resolution maps computed using cryoSPARC. The 0.143 cutoff is indicated with an horizontal blue line in panels c and f. NUR: non-uniform refinement.

**Figure S3.**
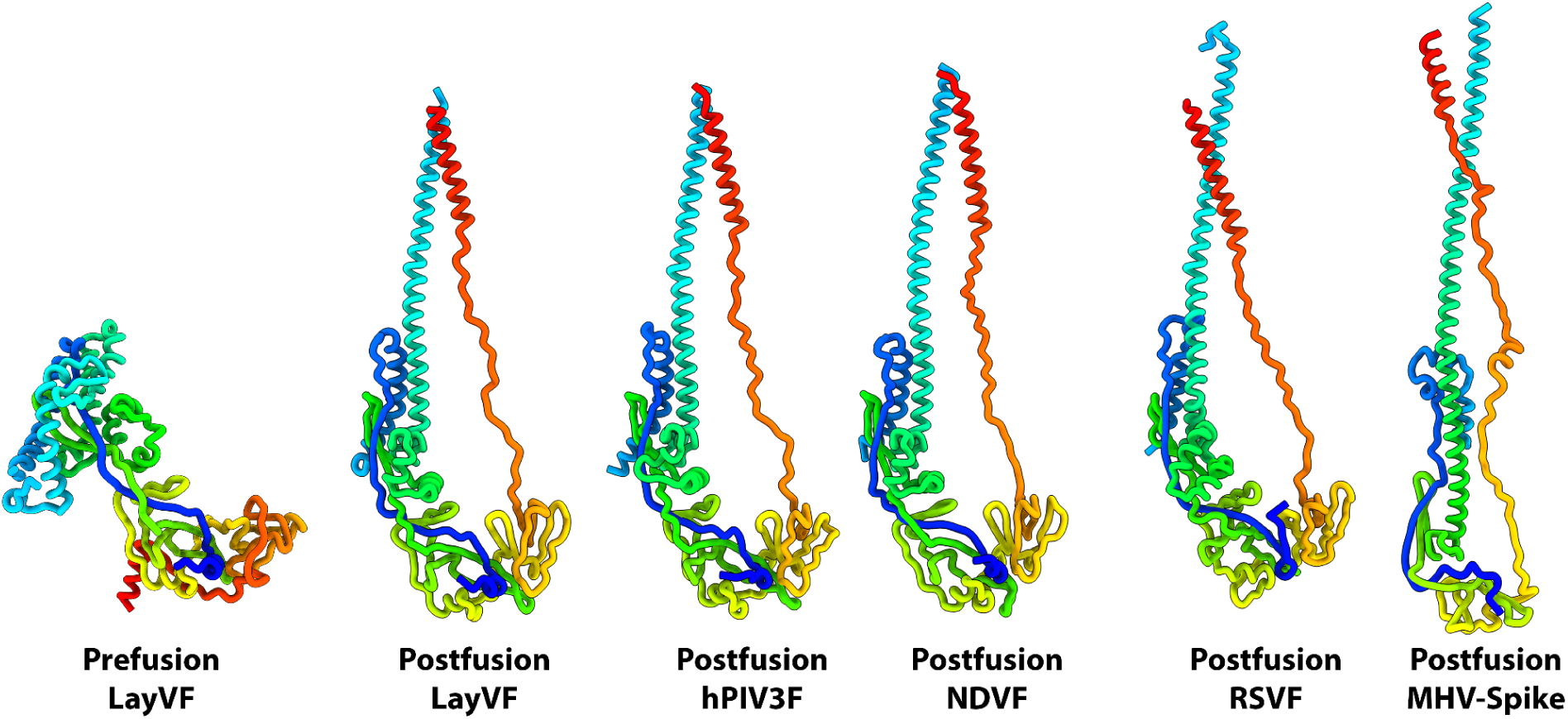
Conservation of the general architecture of paramyxovirus and coronavirus fusion proteins. Ribbon diagrams of viral class I fusion proteins underscoring the architectural conservation among paramyxovirus and coronavirus postfusion structures. All models are colored using a rainbow scheme from blue (N-terminus) to red (C-terminus). The structures rendered are hPIV3 F: PDB 1ZTM; NDV F: PDB 3MAW; RSV F (PDB 3RRR); MHV S (PDB 6B3O).

**Extended Data Fig. 4.**
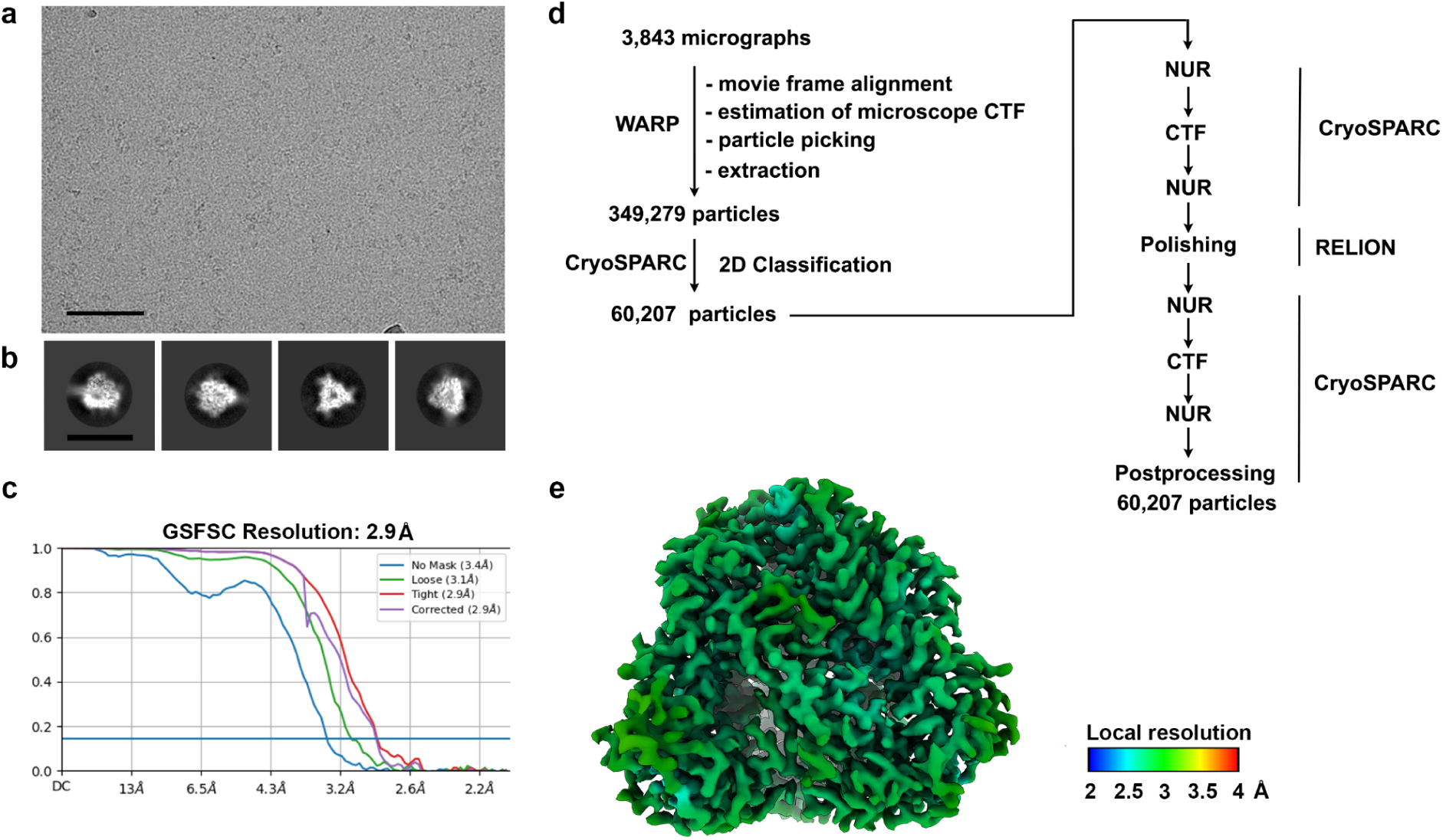
CryoEM data processing pipeline for GhV F. **a-b,** Representative electron micrograph (a) and 2D class averages (b) of GhV F embedded in vitreous ice. The scale bar represents 100 nm (a) or 150 Å (b). **c,** Gold-standard Fourier shell correlation curve for the GhV F reconstruction. The 0.143 cutoff is indicated with an horizontal blue line. **d,** Data processing flowchart. CTF: contrast transfer function; NUR: non-uniform refinement; Polishing: Bayesian particle polishing implemented in Relion. **e,** Local resolution map calculated using CryoSPARC and plotted onto the sharpened GhV F reconstruction.

**Extended Data Fig. 5.**
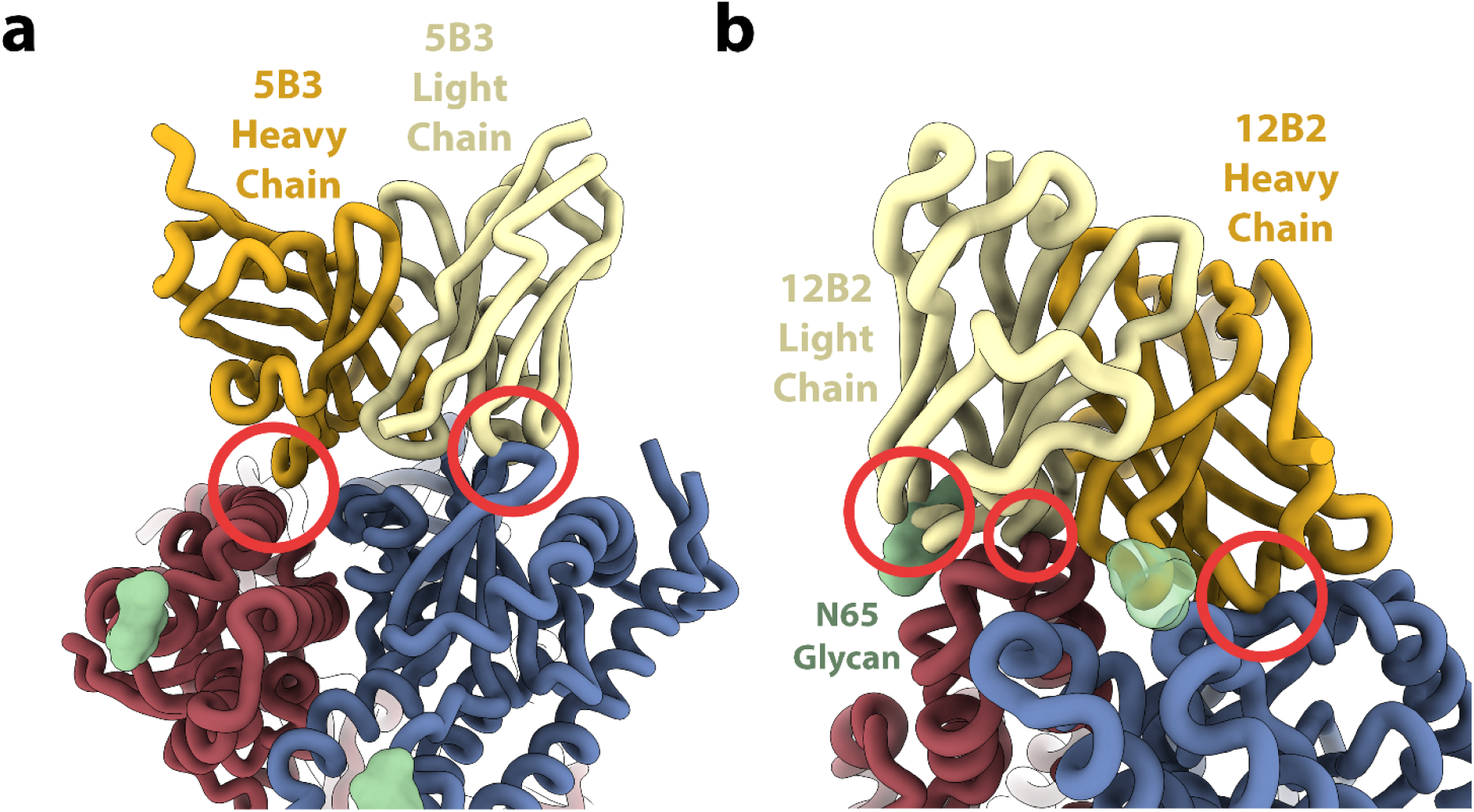
Incompatibility of prefusion LayV F with binding of two NiV/HeV F neutralizing mAbs. **a,** Superimposition of the 5B3-bound NiV F structure (PDB 6TYS) onto prefusion LayV F. Two LayV F protomers are shown in blue and red whereas the 5B3 heavy and light chains are rendered gold and yellow respectively. NiV F is omitted for clarity. N-linked glycans are shown as green surfaces. Red circles indicate potential clashes. **b,** Superimposition of the 12B2-bound HeV structure (PDB 7KI4) onto prefusion LayV F with the same color and representation scheme as in panel a.

**Extended Data Fig. 6.**
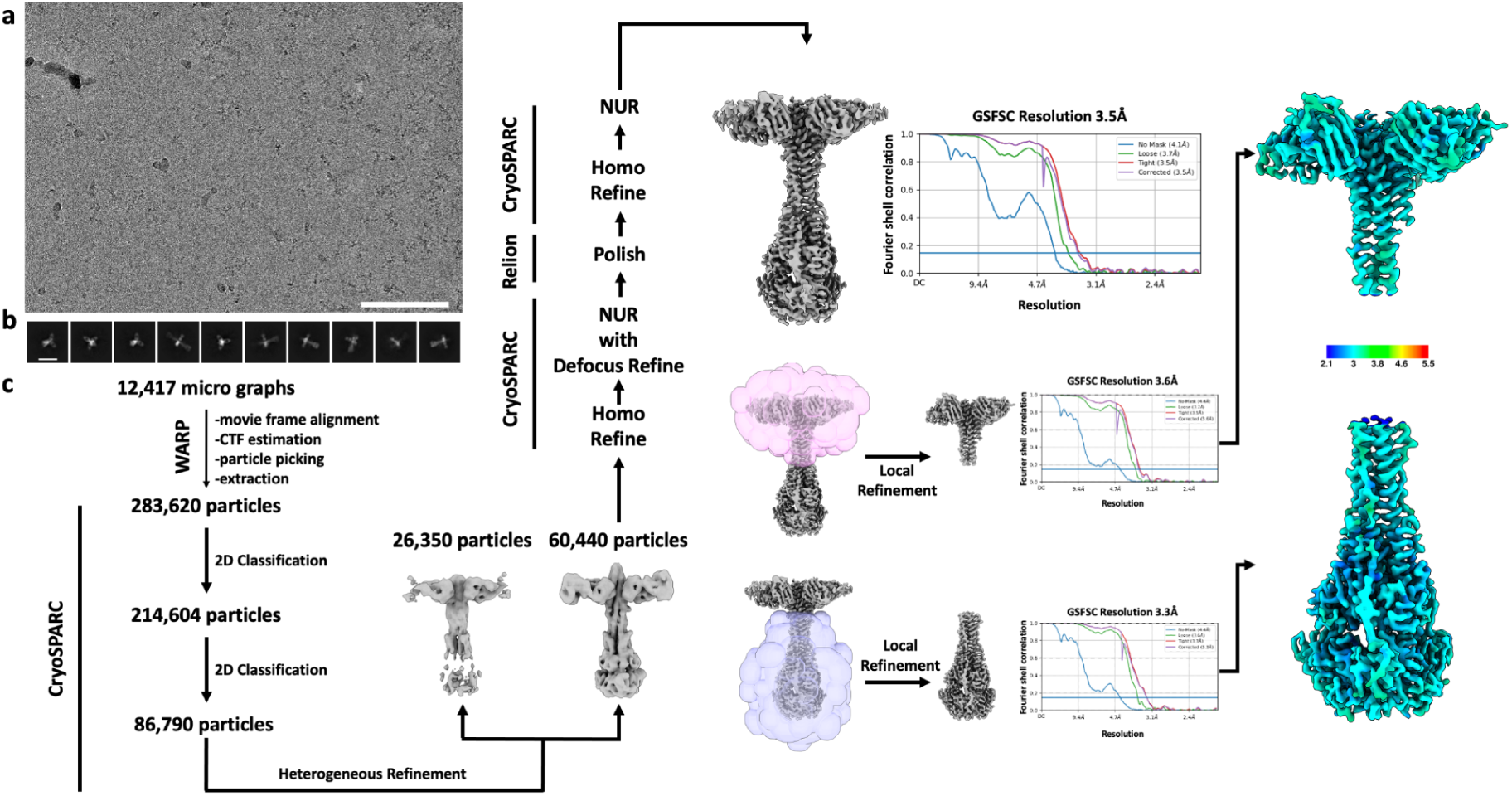
CryoEM data processing pipeline for postfusion LayV F in complex with the 4G5 Fab. **a-b,** Representative electron micrograph (a) and 2D class averages (b) of 4G5-bound postfusion LayV F embedded in vitreous ice. The scale bar represents 100 nm (a) or 200 Å (b). **c,** CryoEM data processing flow chart including local resolution maps computed using cryoSPARC.

**Extended Data Fig. 7.**
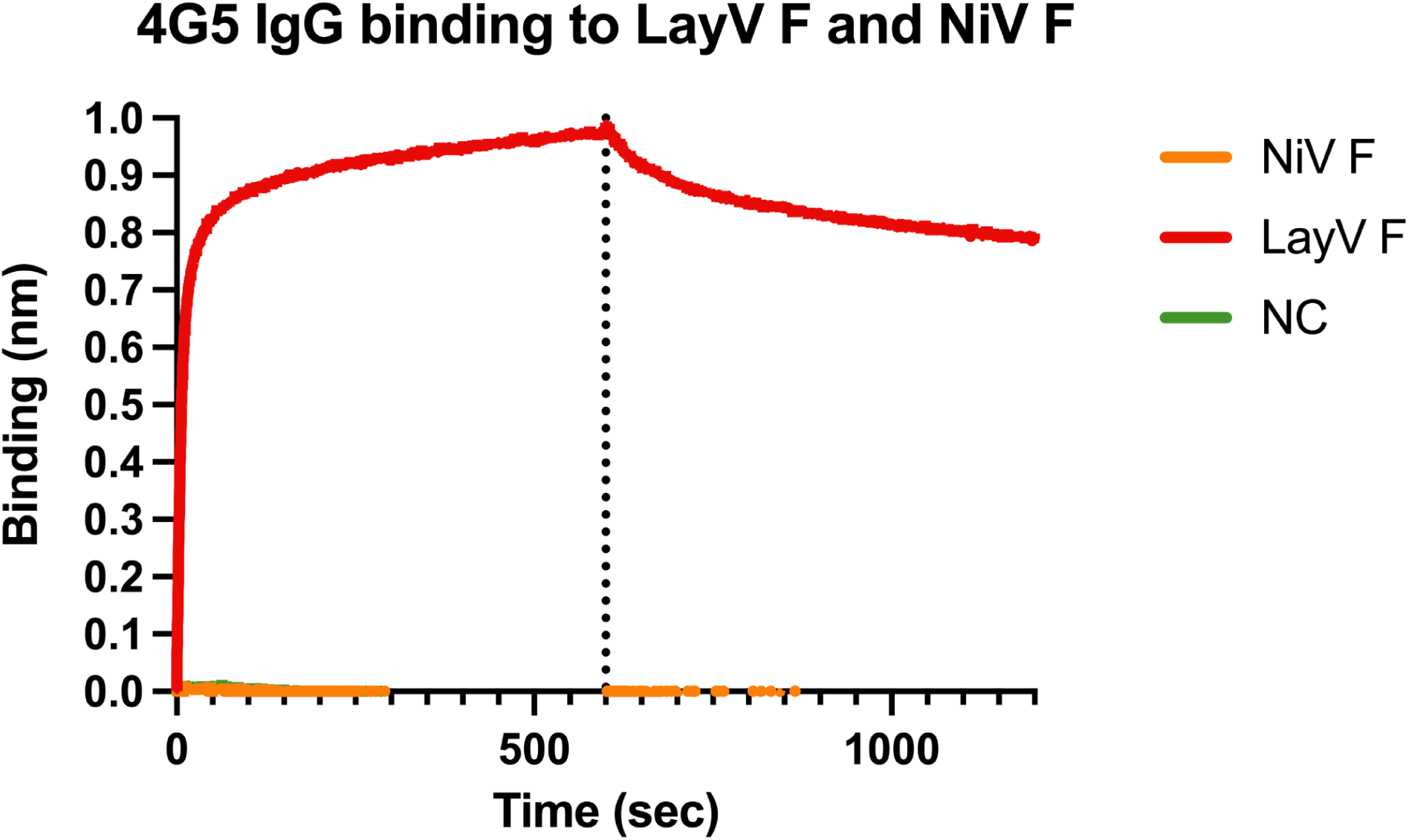
4G5 IgG recognizes LayV F but not NiV F. Biolayer interferometry binding analysis of 100nM of LayV F (red), NiV F (orange) or kinetic buffer NC: negative control, green) to the 4G5 IgG binding immobilized at the surface of AMC biosensors.

**Extended Data Fig. 8.**
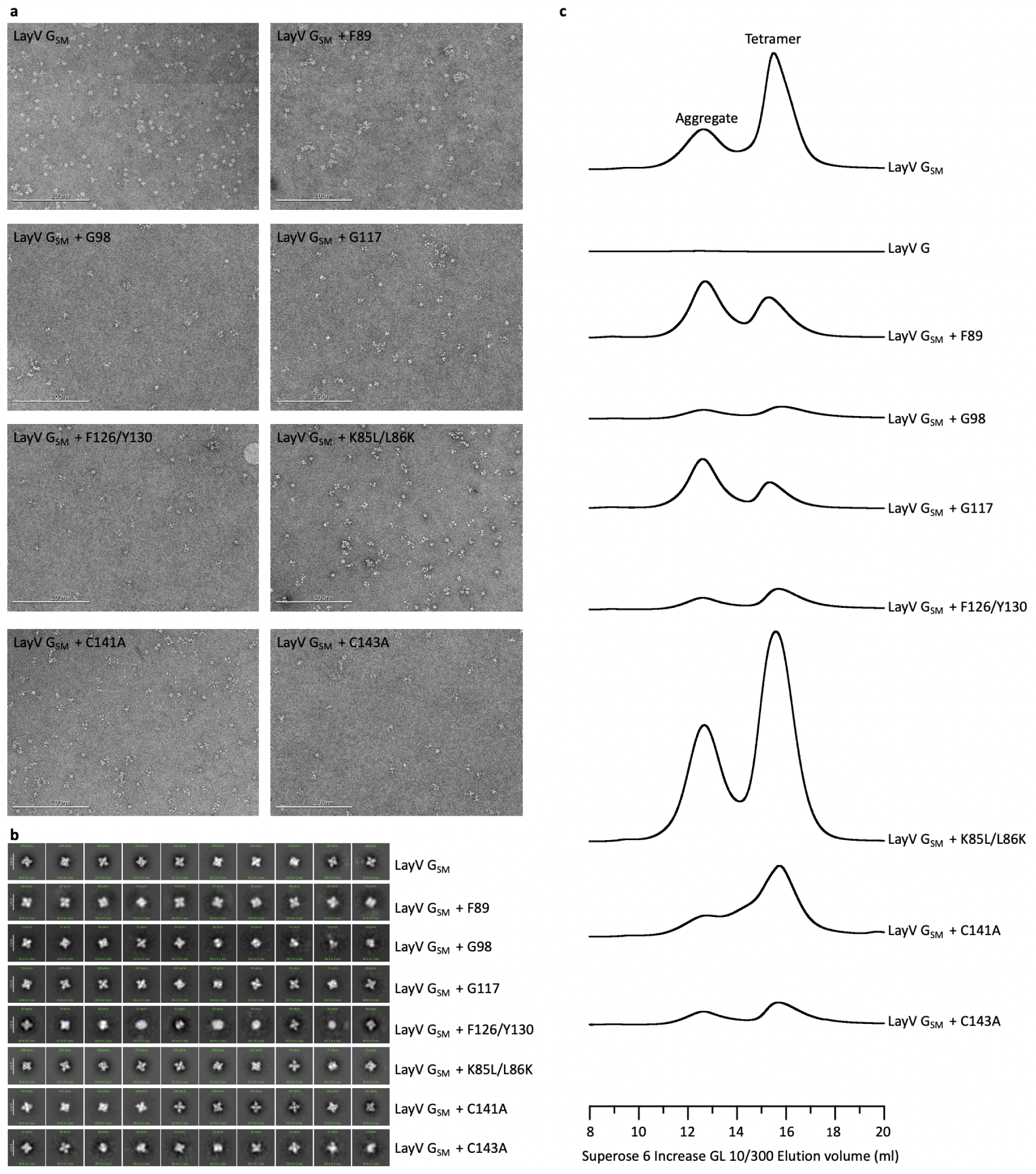
Characterization of designed LayV G mutants. **a**, Electron micrographs of negatively stained LayV G mutants after affinity purification and prior to size exclusion chromatography. **b**, 2D class averages obtained from negatively stained LayV G mutants showing the formation of tetramers for all mutants evaluated. Template picking and a prior round of 2D classification was used to enrich well-folded particles. **c**, Size exclusion chromatography profiles of LayV G mutants highlighting variations in protein aggregation and tetramerization.

**Extended Data Fig. 9.**
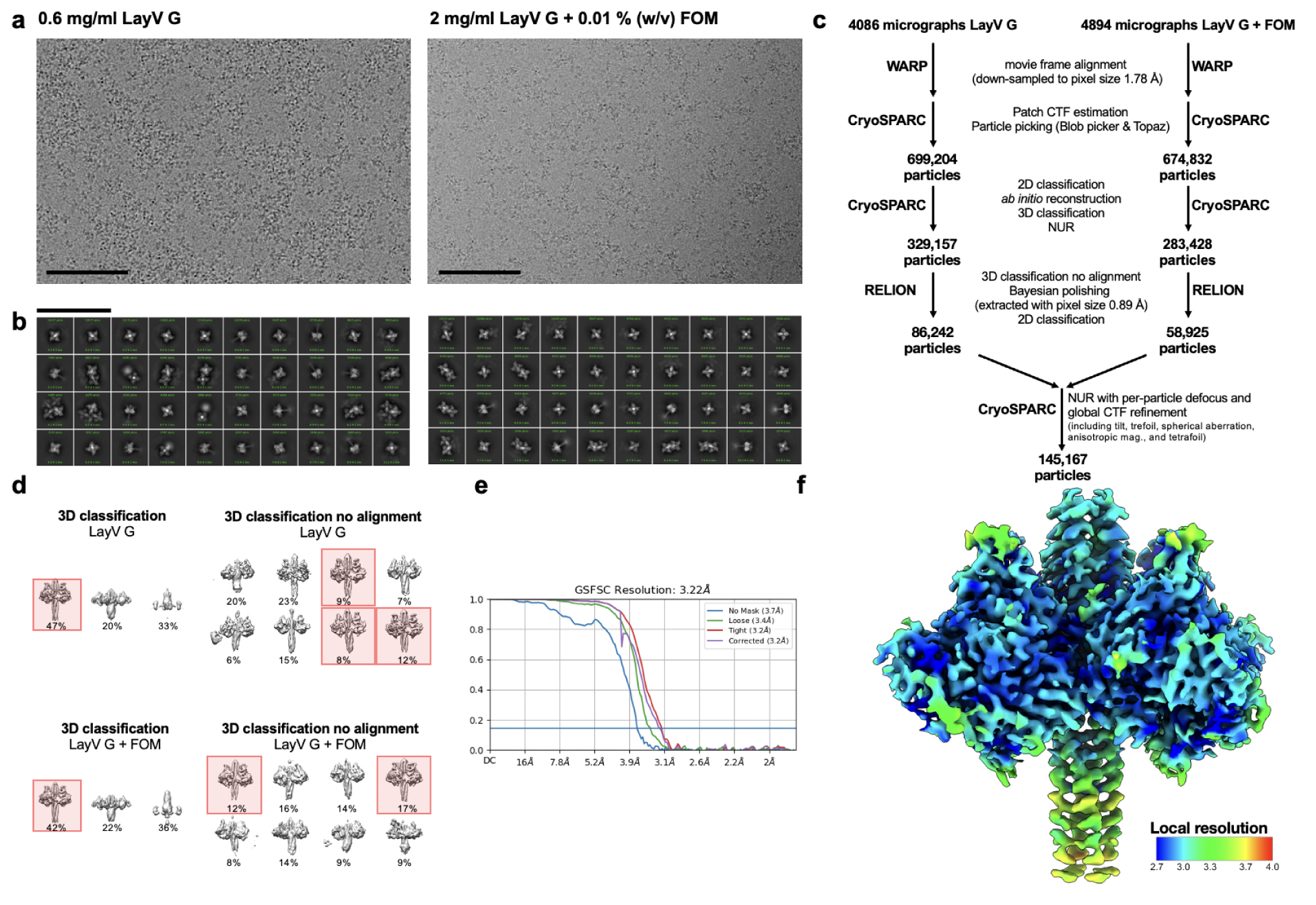
CryoEM data processing pipeline for LayV G_SM_. **a-b**, Representative electron micrograph (a) and 2D class averages (b) of LayV G_SM_ embedded in vitreous ice. The scale bars represent 100 nm. **c**, Data processing flowchart. CTF: contrast transfer function; NUR: non-uniform refinement. **d**, 3D maps corresponding to 3D classifications with and without alignment referenced in the data processing flow chart. Classes highlighted in red were used for further data processing. **e**, Gold-standard Fourier shell correlation curve for the LayV G reconstruction. The 0.143 cutoff is indicated by the blue line. **f**, Local resolution map calculated using CryoSPARC and plotted onto the sharpened LayV G reconstruction.

**Extended Data Fig. 10.**
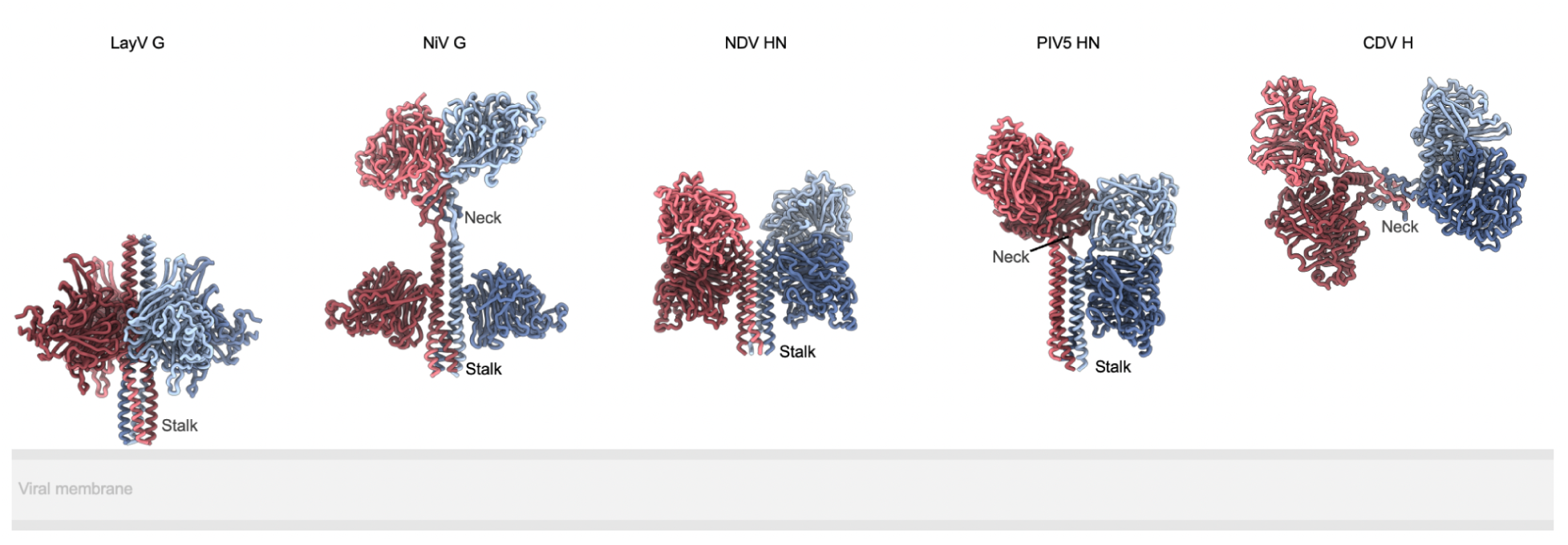
Comparison of paramyxovirus attachment glycoprotein architectures. Ribbon diagrams of LayV G, NiV G (PDB 7TY0 and 7TXZ), NDV HN (PDB 3T1E), PIV5 HN (PDB 4JF7) and CDV H (PDB 7ZNY) aligned by their stalks to show relative distance from and orientation to the viral membrane.

**Extended Data Fig. 11:**
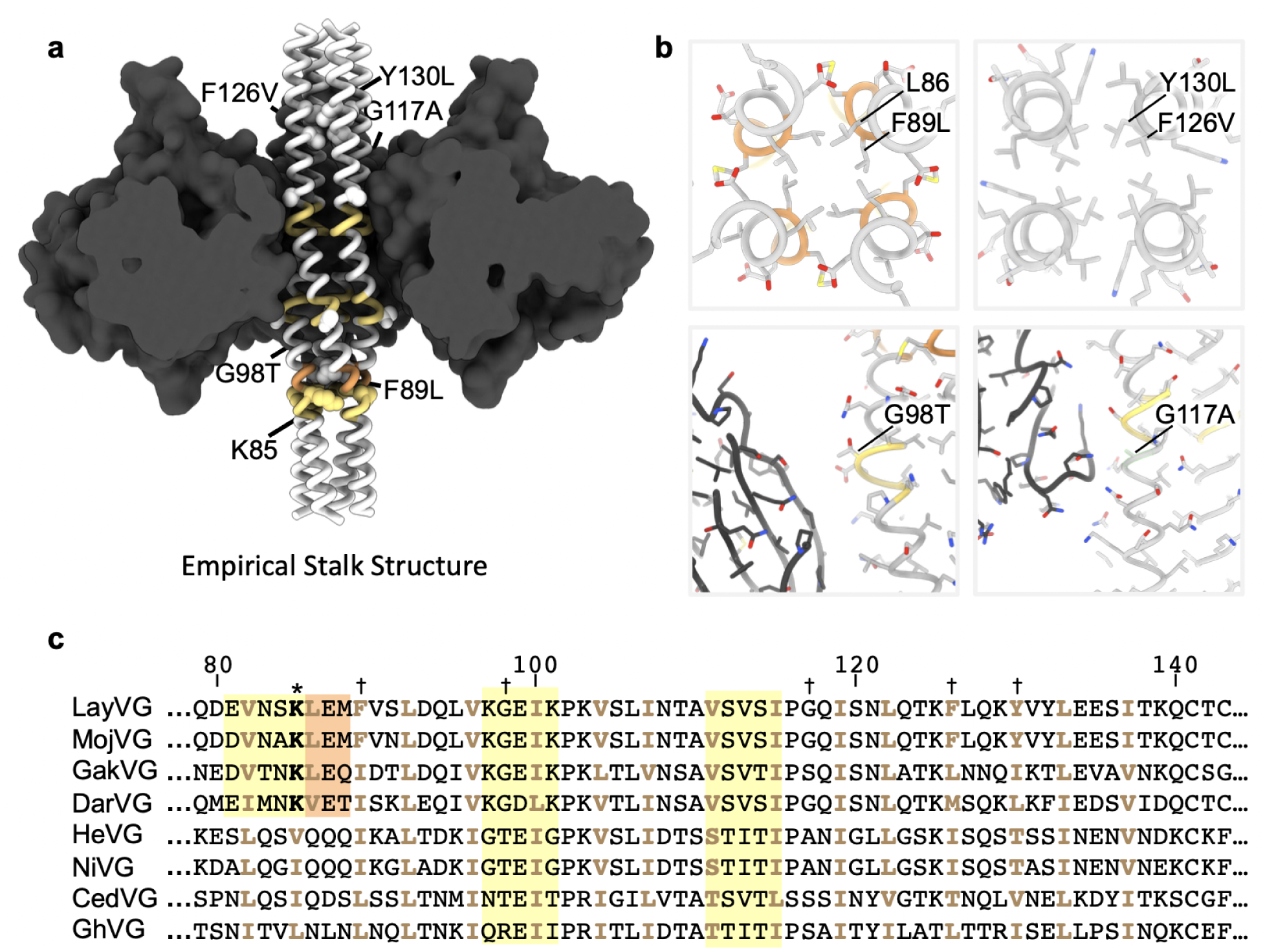
Stalk stabilizing mutations and comparison of the stalk sequence with other HNV G glycoproteins. **a,** Designed LayV G stabilizing mutations (labeled white spheres) in the stalk of the LayV G_SM_ cryoEM structure highlighting pi-helices (yellow), and 3_10_-helices (orange). The head domains are shown as black surfaces. **b**, Zoomed-in views of stabilizing mutations with relevant and neighboring side chains rendered as sticks colored as in (a). **c,** Amino acid sequence alignment spanning LayV G residues 79-143 with other HNV G glycoproteins. Known or predicted residues facing the interior of the tetrameric coiled coil are shown in beige. Regions with known or predicted pi-helical structure are highlighted in yellow. Regions with known or predicted 3_10_-helical structure are highlighted in orange. K85 is marked with an asterisk. Residues mutated for stability are indicated with a dagger.

**Extended Data video 1.** 3DFlex cryoEM density series oscillating between 41 frames showing movement of the LayV G stalk.

## Notes

### Summary of Updates

Title change

